# A novel vaccine targeting the viral protease cleavage sites protects Mauritian cynomolgus macaques against vaginal SIVmac251 infection

**DOI:** 10.1101/842955

**Authors:** Hongzhao Li, Robert W. Omange, Binhua Liang, Nikki Toledo, Yan Hai, Lewis R. Liu, Dane Schalk, Jose Crecente-Campo, Tamara G. Dacoba, Andrew B. Lambe, So-Yon Lim, Lin Li, Mohammad Abul Kashem, Yanmin Wan, Jorge F. Correia-Pinto, Xiao Qing Liu, Robert F. Balshaw, Qingsheng Li, Nancy Schultz-Darken, Maria J. Alonso, James B. Whitney, Francis A. Plummer, Ma Luo

## Abstract

After over three decades of research, an effective anti-HIV vaccine remains elusive. Unconventional and novel vaccine strategies are needed. Here, we report that a vaccine focusing the immune response on the sequences surrounding the 12 viral protease cleavage sites (PCSs) provides greater than 80% protection of Mauritian cynomolgus macaques (MCMs) against repeated intravaginal SIVmac251 challenges. The PCS-specific T cell responses are correlated with vaccine efficacy. The PCS vaccine does not induce immune activation and inflammation known to be associated with increased susceptibility to HIV infection. Machine learning analyses revealed that the immune environment generated by the PCS vaccine predicts vaccine efficacy. Our study demonstrates for the first time that a novel vaccine which targets viral maturation, but lacks full Env and Gag proteins as immunogens, can prevent intravaginal infection in a highly stringent NHP/SIV challenge model. Targeting HIV maturation thus offers a novel approach to developing an effective HIV vaccine.

**One Sentence Summary:** The anti-PCS T cell responses and the immune environment induced by the novel PCS vaccine are key correlates of vaccine efficacy

## Introduction

HIV mutates rapidly and preferentially infects activated CD4+ T cells. Inflammation activates and attracts HIV target cells, thereby enhancing the risk of infection^1-3^. These inherent characteristics of HIV infection underscore the greater challenges compared to other pathogens when developing a prophylactic vaccine ^4,5^. Among the six HIV vaccine clinical efficacy trials to date, only the RV144 Thai trial demonstrated any efficacy, and it was modest (31.2%)^4,6^. Since activated CD4 T cells are preferential targets for HIV-1, an effective HIV vaccine would need to elicit protective anti-HIV immunity while minimizing vaccine-induced inflammation and immune activation^5,7,8^. Our previous studies of HIV highly exposed seronegative (HESN) Kenyan female sex workers showed that narrowly targeted T cell responses were associated with protection against HIV infection^8-10^. Thus, an alternative strategy for an effective HIV vaccine would be to generate immune response only to the essential part of HIV-1^8,11^. Learning from natural immunity observed in these Kenyan HESN women, we tested a novel strategy by focusing the host immune response on the sequences surrounding the 12 viral protease cleavage sites (PCSs) (the PCS vaccine)^7,12^. The HIV protease cleaves Gag, Gag-Pol and Nef precursor proteins at the 12 PCSs^7,13^. Studies have shown that the process of protease cleavage requires a tightly controlled and ordered sequence of proteolytic processing events mediated by different rates of cleavage at the different processing sites^14-20^. Even subtle disturbances may be sufficient to interrupt this delicately balanced process and drive it toward a non-productive end^14,17,18,21^. We hypothesized that a vaccine targeting the 12 PCSs would be effective against HIV infection. First, as the sequences surrounding the PCSs are highly conserved^7^ the vaccine can target multiple HIV subtypes. Second, the focused immune response may drive viral mutations surrounding the PCSs and impair the proteolytic processing of HIV polyproteins. As a result, non-infectious viral progenies will be generated. Third, restricting the immune response to a smaller number of T-cell epitopes surrounding the PCS may result in lower vaccine-induced inflammation and fewer activated CD4 T target cells^7^. We tested this hypothesis in a non-human primate (NHP)/simian immunodeficiency virus (SIV) infection model. We demonstrate here that the PCS vaccine, without full Gag and Env immunogens, protects female Mauritian Cynomolgus macaques (MCMs)^22^ from acquiring highly pathogenic and heterogenic SIVmac251^23^ infections.

## Results

### Vaccine protection against vaginal SIVmac251 infection

The PCS vaccine delivers twelve 20-amino acid peptides overlapping each of the 12 viral protease cleavage sites with recombinant vesicular stomatitis viruses (rVSV) and nanoformulations (NANO) ^7,12^. To evaluate the efficacy of the PCS vaccine (Figure 1A), 16 female MCMs that did not express known SIV controller MHC haplotypes^24^ (Table S1) were divided into two groups (the PCS vaccine and the vaccine vector control; n=8 per group). MHC haplotypes were balanced between the two groups (Table S2). The vaccination scheme consisted of prime with rVSVpcs or rVSV control vector (intramuscularly) and four boosts with combinations of rVSVpcs or rVSV vector (intramuscularly) and NANOpcs (intranasally)^12^ (Figure 1B).

**Figure 1.**
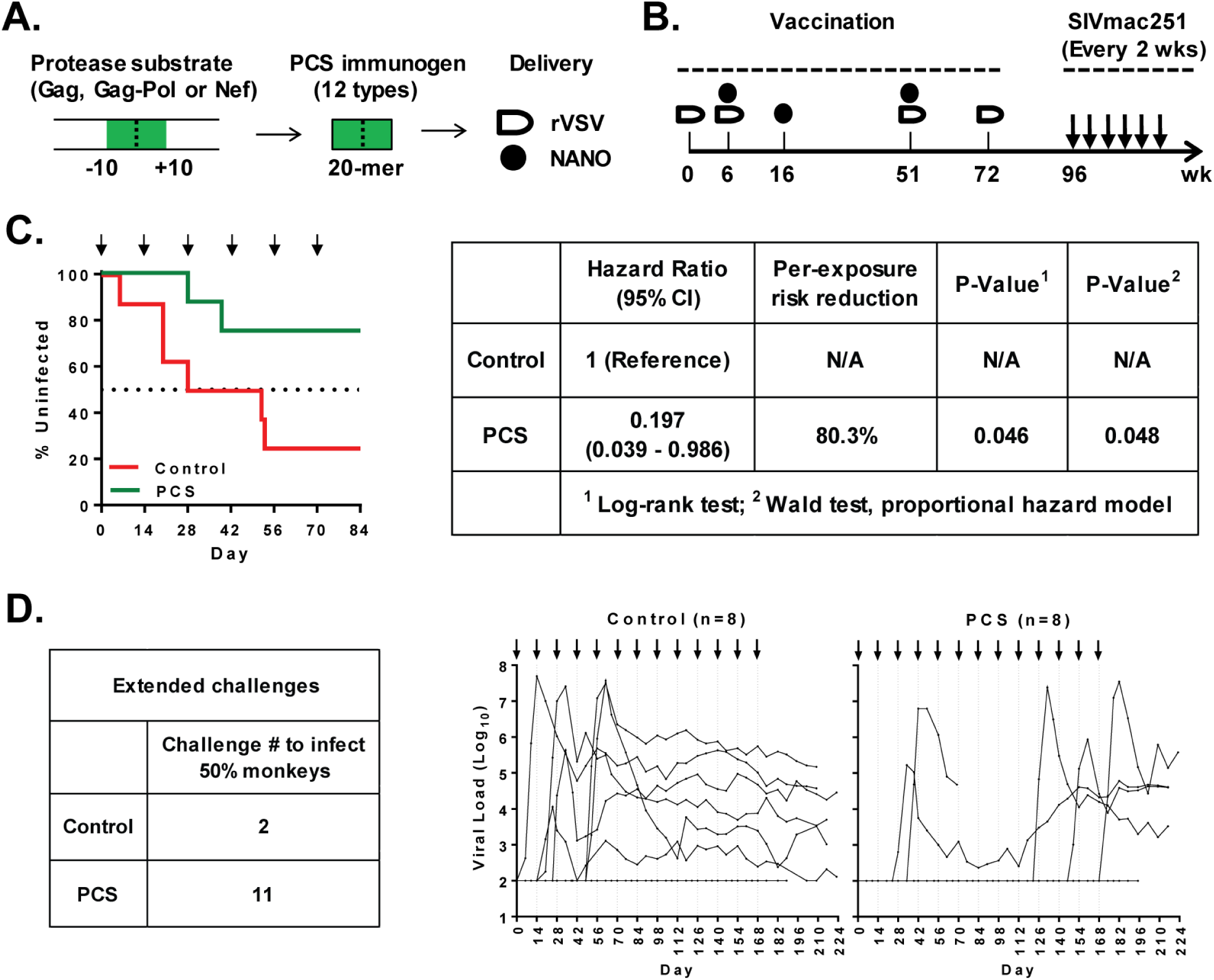
The PCS vaccine protected MCMs against vaginal SIVmac251 infection. (A). The PCS vaccine. Twelve 20-mer peptides derived from the twelve protease cleavage site (PCS) regions of SIVmac239 (between −10 and +10 positions flanking each cleavage site) were delivered as recombinant vesicular stomatitis viruses (rVSV) and nanoparticles (NANO). (B). Immunization and challenge scheme. Two groups of animals were involved, the PCS vaccine group (n = 8) and the sham vaccine control group (n = 8). (C) Vaccine efficacy. Left: Kaplan-Meier plot showing the percentage of uninfected animals following challenges. Right: Statistical analyses of vaccine efficacy. (D). Extended challenges beyond the pre-determined, standard challenge protocol. Left: Numbers of challenges required to infect 50% animals in each group. Right: Viral load time course of each animal (note: n = 8 per group while data of some animals at baseline levels overlap and are not visually distinguishable on graph).

Approximately six months after the final boost, repeated low-dose intravaginal SIVmac251 challenges (the Desrosiers 2010-Day 8 stock at 250 TCID50/animal/challenge) were carried out every two weeks and infection status was monitored by quantification of plasma viral load on days 6, 10 and 14 after each of the challenges (Figure 1B). The vaginal challenge route was chosen to mimic vaginal HIV exposure in women, which accounts for about half of all HIV infections worldwide^25^. According to our study design (detailed in Methods), the end point of the challenge study for evaluating vaccine efficacy was at challenge number six (Figure 1C), which has also been used by many other studies^23,26-28^. At the end of this standard challenge procedure, the majority (six of eight, 75%) of control animals were infected, whereas only two of the PCS-vaccinated animals (25%) were infected (Figure 1C). To determine whether continued challenges can lead to infection of the two uninfected control animals and to determine the numbers of challenges needed to infect 50% of the vaccinated animals because of biological variations among the monkeys, we extend the challenges to thirteen times (Table S3). Following the extended challenges three additional animals in the PCS vaccine group became infected (after 9, 11 and 13 challenges, respectively). However, the two previously uninfected control animals remained uninfected (Table S3). This observation was consistent with findings from several previous studies in rhesus macaques that a proportion of control animals were refractory to acquisition of SIV/SHIV infections^28-36^.

Survival analysis demonstrated that the PCS vaccine significantly increased the number of challenges required for acquisition of SIVmac251 infection (p = 0.046, log-rank test) (Fig. 1C). It provided an 80.3% reduction in the per-exposure risk of viral acquisition (vaccine efficacy, VE = 1 – hazard ratio) (proportional hazards regression) (Figure 1C). Furthermore, while it took only two SIVmac251 challenges to infect 50% animals in the control group (Figure 1C and 1D), extended challenges showed that it took 11 challenges to infect 50% animals in the PCS vaccine group (Figure 1D). The observed vaccine protection was not related to the menstrual phases of these female animals during the SIVmac251 challenges (Figures. S1 and S2) or their MHC haplotypes (Tables S4 and S5).

These data demonstrate for the first time that a candidate prophylactic PCS HIV vaccine, without traditional types of immunogens such as full Gag and Env proteins, significantly protected female monkeys against highly pathogenic and heterogenic SIVmac251 challenges.

### The PCS vaccine did not elicit significant inflammatory responses in the cervico-vaginal mucosa

Inflammation can activate and attract HIV CD4 T target cells to the portal of entry. Elevated cervicovaginal mucosal (CVM) inflammation is associated with increased susceptibility to HIV/SIV infection and reduced efficacy of anti-HIV microbicides^37-39^. We examined vaginal mucosal inflammation following vaccination. The vaccine-induced mucosal inflammation was analyzed by quantifying inflammatory cytokine responses in cervicovaginal lavage (CVL) samples (Figure 2 A-G and Figure S3). Thirteen pro-inflammatory cytokines (TNF-α, IFN-γ, IL-6, RANTES/CCL5, GM-CSF, IL-1β, MCP-1/CCL2, IL-8, MIP-1α/CCL3, MIP-1β/CCL4, IP-10/CXCL10, IL-17A and IL-1α) and one anti-inflammatory cytokine (IL-10) were analyzed. Vaccine-induced fold changes in cytokine levels at multiple time points were determined and shown as heat maps (Figure 2A). No apparent induction of inflammatory cytokine responses, apart from some sporadic fluctuations, was observed in the control or the PCS vaccine group. The short SIV PCS peptide immunogens in the PCS vaccine did not lead to the persistent increase in cervicovaginal inflammatory responses in the PCS vaccine group in comparison to the control group, except an initial increase after prime in IFN-γ, IL-6, RANTES, GM-CSF, MCP-1, and IL-17A, which were dampened at later boost points (Figure 2A). When the absolute cytokine levels were compared, the PCS vaccine group had significantly lower levels of IFN-γ and MIP-1β at week 73 (one week after the final boost), and IL-8 and MIP-1β at week 90 (18 weeks after the final boost), than that of the control group (Figure 2B-G). These data indicate that the PCS vaccine did not elicit significant inflammatory responses in the cervico-vaginal mucosa.

**Figure 2.**
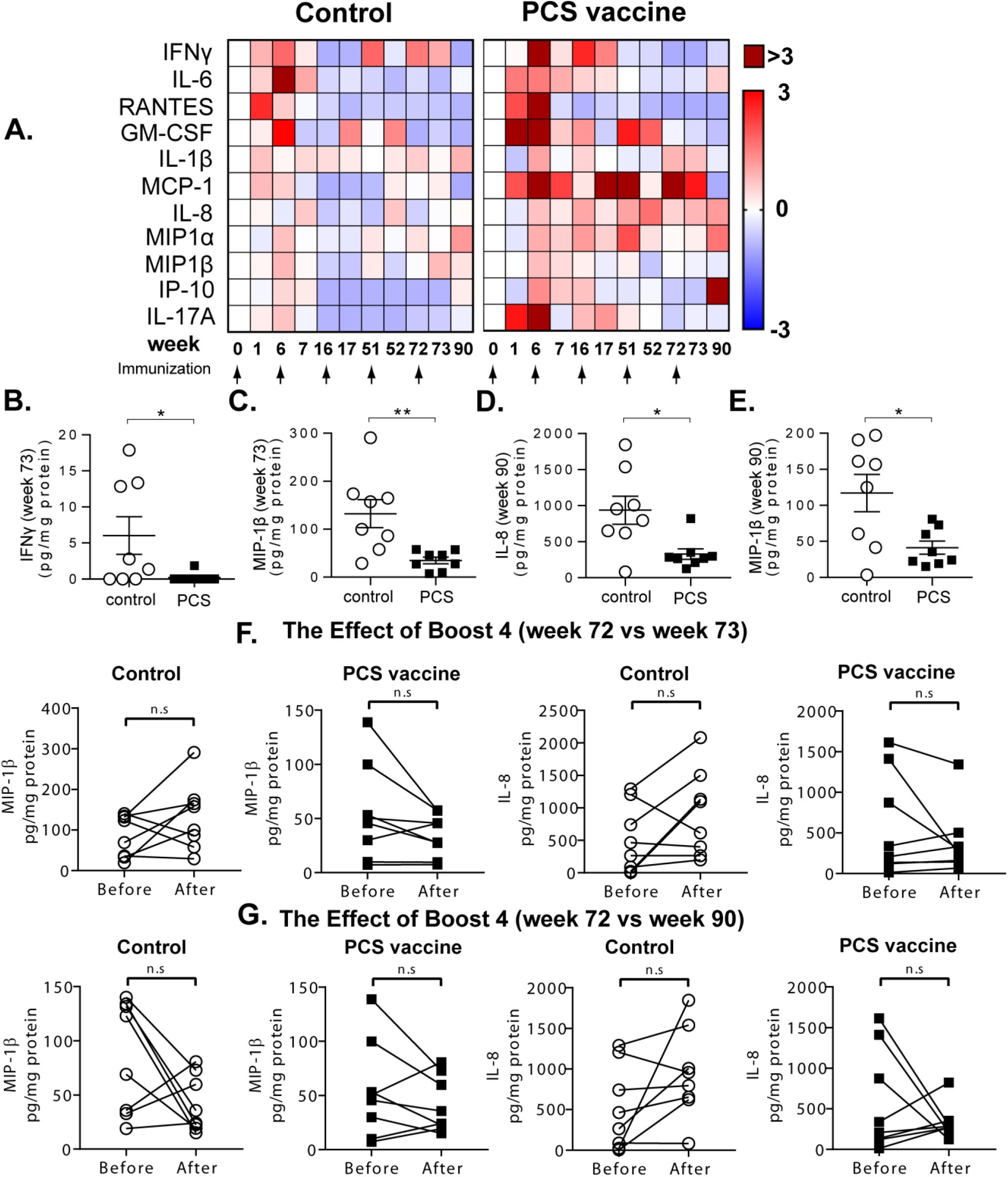
The PCS vaccine and mucosal inflammatory cytokines. (A). Heat maps showing net fold change (relative to pre-immunization baseline) in group median inflammatory cytokine levels at the cervicovaginal mucosa at different time points of the vaccination scheme. (B-E). The PCS vaccine group showed significantly lower levels of several cytokines at indicated time points than the control group. Data are presented as values from individual animals with mean ± SEM. *p < 0.05 and **p < 0.01 (Mann-Whitney’s test). (F). Comparisons of the cytokines in B-E at 4th boost (week 72) and a week after (week 73). (G). Comparisons of the cytokines in B-E at 4th boost (week 72) and 18 weeks after (week 90).

### The PCS vaccine induced antibody and cellular responses

Mucosal antibodies to PCS peptide antigens were measured in CVL (reported previously)^12^ and plasma samples (Figure S4A and Fig. S5). The patterns of anti-PCS peptide antibody (IgG) responses in CVL and plasma were similar. A notable trend of increase in antibodies to individual PCS antigens was observed at week 73 (one week after the last boost, defined as the “peak” immune response time point) (Fig. S5). The level of combined plasma antibodies to all 12 PCS peptides at this time point in the PCS vaccine group was approximately two fold of that in the control group (Figure S4A). Thus, the PCS vaccine induced antibodies to PCS peptides. The PCS peptides are not located in the Env, and the pre-challenge plasma samples of the PCS vaccine group did not have neutralizing activities against tier 1 and tier 2 SIVmac251 (data not shown). The moderate magnitude of anti-PCS antibodies may reflect the existence of B cell epitopes in the sequences surrounding the PCS, and indicate that immunization with the PCS vaccine was effective in inducing antibodies to the PCS peptides. However, the contribution of these antibodies to the protection is currently unknown.

To examine cellular immune responses, peripheral blood mononuclear cells (PBMCs) from the “peak” time point were quantified by ELISPOT for frequency of IFNγ-producing cells in response to a pool of SIV peptide antigens (PCS1-4, PCS5-8 or PCS9-12) (Figure S4 B-D). Slightly higher numbers of animals in the PCS vaccine group than those in the control group showed above background responses to PCS5-8 and PCS9-12 peptide pools, but these differences were not statistically significant (Figure S4 B-D). Although IFNγ ELISPOT has been traditionally used to evaluate T cell responses to vaccines, many cytokines other than IFNγ are known to be secreted by activated T cells^40-42^. Thus, simply assessing the IFNγ response by ELISPOT may give a very limited view of vaccine-induced T cell responses^43^. Furthermore, in some vaccine studies, IFNγ secretion was a poor correlate of protection against HIV, and was not the best indicator of vaccine-induced responses^43^. We therefore expanded the analysis of cellular responses by measuring 14 cytokines using our customized Bio-Plex multiplexed cytokine assay.

PBMCs of the PCS vaccine group from the “peak” time point were stimulated with PCS peptide pools and quantified for cytokines secreted into the culture supernatants (Table S6). Between nine to fourteen cytokines were detected following stimulations with different PCS peptide pools (PCS1-4: nine cytokines, PCS5-8 and PCS9-12: fourteen cytokines). Among these cytokines, IFNγ was at a low level and detectable only in less than half of the animals. Six other cytokines, including RANTES, MIP-1α, MIP-1β, TNF-α, IL-6 and IL-8, were detected in 50% or higher percentage of animals (Table S6). Some of these cytokines are currently known to have potential functions in directly inhibiting HIV infection or regulating inflammation. RANTES, MIP-1α and MIP-1β are HIV-suppressive factors produced by CD8^+^ T cells possibly through competitive binding of HIV co-receptor CCR5^44^. IL-6, produced by various cell types including T cells, B cells and monocytes^45^, can have an anti-inflammatory role in controlling local or systemic acute inflammatory responses^46,47^. A source of IL-8 is CD4^+^FOXP3^+^ regulatory T cells^48^, which could play a role in regulating antiviral response. In summary, these data indicated that several types of cellular immune responses to PCS antigens can be generated following vaccination.

### The PCS vaccine generated fewer viral target cells and higher PCS peptide specific CD8^+^ T memory and CD4^+^ T_reg_ responses

We characterized the T cell subsets after immunization with the PCS vaccine. PBMCs from week 90 (a pre-challenge time point) without antigen stimulation (*ex vivo*) or stimulated with a pool of PCS peptides (PCS1-12) were analyzed by flow cytometry (Figure 3, 4 and Figure S6). Different cellular markers^49-51^ were chosen to differentiate CD4^+^ and CD8^+^ T cell subsets including naive and memory T cells (CCR7 and CD45RA)^52,53^, Th17 cells^4,54^ (IL-17A and CCR5) and Treg cells (CD25, FoxP3 and CD127)^54^, together with activation and differentiation markers including CD38, CD69, CD107a, Ki-67, PD-1, IL-2, TNF-α, IFN-γ, MIP-1β and IL-10.

**Figure 3.**
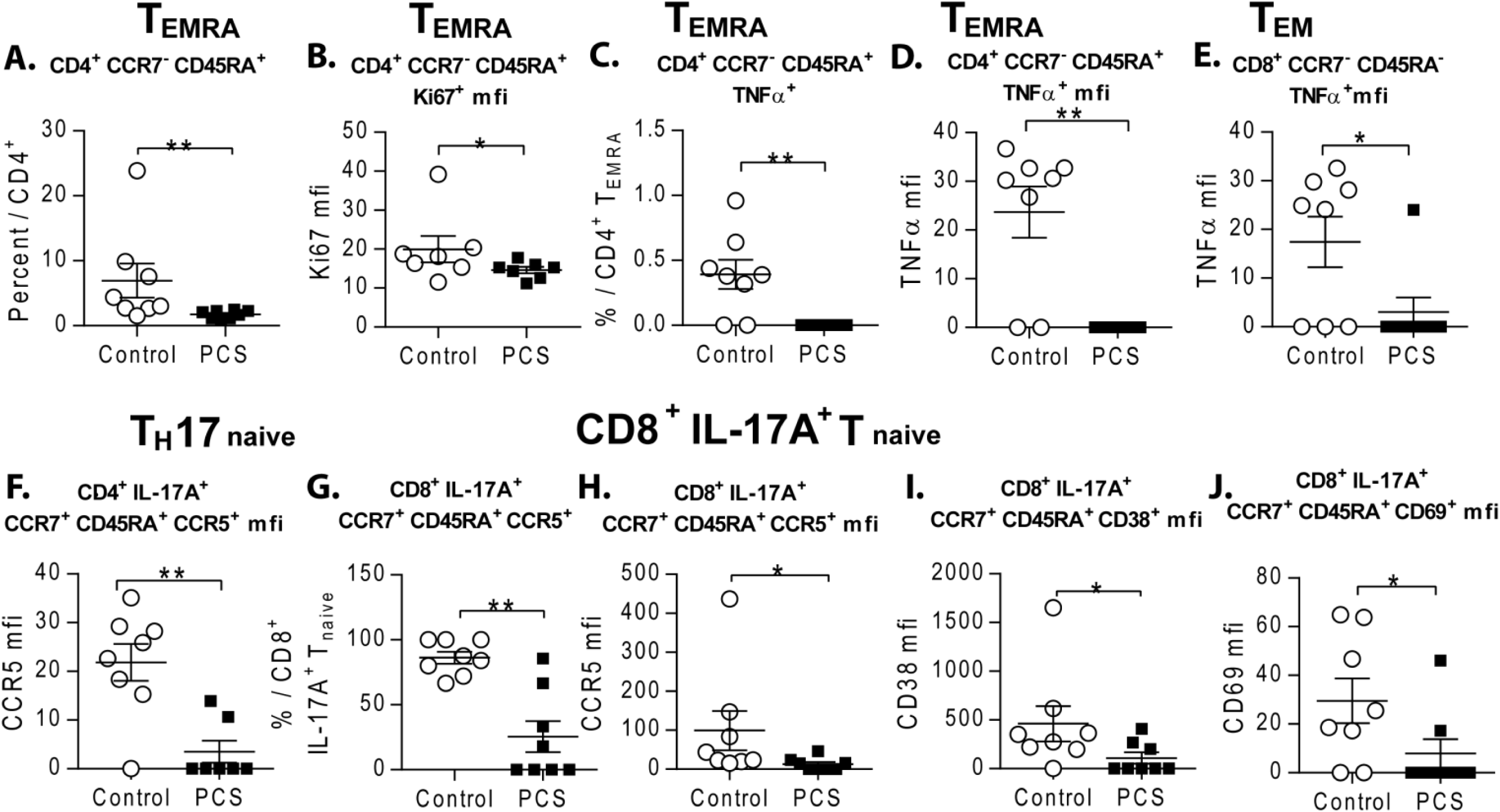
The PCS vaccine induced CD4 and CD8 T cell responses (Ex vivo). PBMCs from the pre-challenge time point (week 90) without antigen stimulation (ex vivo) were analyzed by multicolor flow cytometry. Definition of T cell subsets based on cellular markers. TCM: central memory T cells, T naive: naïve T cells, TEM: effector memory T cells and TEMRA: CD45RA+ effector memory T cells. Gating strategy is shown in Fig. S6. A-J: Comparisons of cellular subsets between the control and PCS vaccine group. Data are presented as values from individual animals with median lines. Statistical significance of difference was determined by Mann-Whitney’s test: *p < 0.05.

**Figure 4.**
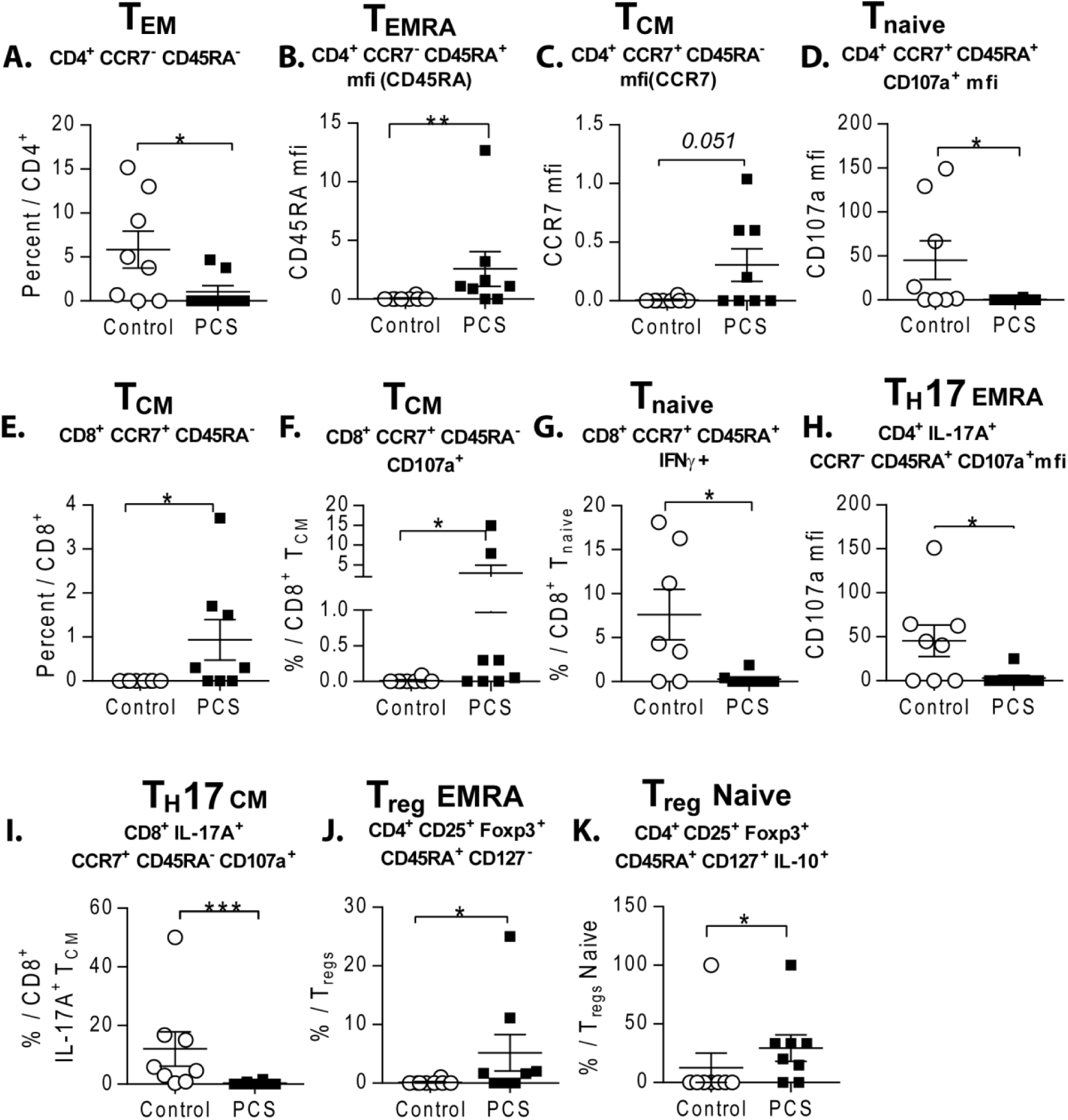
The PCS vaccine induced CD4 and CD8 T cell responses (recall). PBMCs from the pre-challenge time point (week 90) stimulated with antigen (a total PCS peptide pool:PCS1-12), were analyzed by multicolor flow cytometry. Definition of T cell subsets based on cellular markers. TCM: central memory T cells, Tnaive: naïve T cells, TEM: effector memory T cells and TEMRA: CD45RA+ effector memory T cells. Gating strategy is shown in Fig. S6. A-K: Comparisons of cellular subsets between the control and PCS vaccine group. Data are presented as values from individual animals with median lines. Statistical significance of difference was determined by Mann-Whitney’s test: *p < 0.05.

Under *Ex vivo* conditions, compared to the control group the monkeys of the PCS vaccine group had lower frequency of CD4^+^CCR7^−^CD45RA^+^ terminal effector CD4 T cells (CD4^+^ T_EMRA_) and lower frequency of TNF-α+CD4^+^ T_EMRA_ cells. These CD4^+^ T_EMRA_ cells expressed lower amounts of proliferation marker Ki67 and pro-inflammatory cytokine TNF-α (Figure 3A-D). Similarly, CD8^+^CCR7^−^ CD45RA^+^ terminal effector CD8 T cells (CD8^+^ T_EMRA_) of the PCS vaccine group expressed lower levels of TNF-α (Figure 3E). These suggest that the monkeys of the PCS vaccine group had an overall lower level of immune activation.

After PCS peptide stimulation the monkeys in the PCS vaccine group had a lower frequency of CD4^+^CCR7^−^CD45RA^+^ terminal effector CD4 T cells (CD4^+^ T_EMRA_) cells (Figure 4A). While the surface expression of CD45RA on CD4^+^ T_EMRA_, and CCR7 expression on CD4^+^CCR7^+^CD45RA^−^ T central memory (CD4^+^ T_CM_) were significantly higher in the PCS vaccine group (Figure 4B, 4C). The PCS-vaccinated animals also had lower CD107a expression in naive CD4+CCR7+CD45RA+ T cells (CD4+ Tnaive) (Figure 4D), but significantly higher frequency of CD8^+^CCR7^+^CD45RA^−^ T central memory cells (CD8^+^ T_CM_) (Figure 4E) and higher proportions of them expressing CD107a (Figure 4F). However, the PCS-vaccinated monkeys had fewer CD8^+^ T_CM_ cells expressing IFN-γ^+^ (Figure 4G).

Thus, the frequency of PCS specific CD4^+^ T_EM_ was low, but the surface expression of CD45RA on CD4^+^ T_EMRA_ cells and CCR7 on CD4^+^ T_CM_ cells were higher in the PCS vaccine group. Consistent with lower immune activation status, the PCS peptide specific CD4+ T naive cells of the PCS vaccine group had lower level of CD107a expression. The higher frequency and higher expression level of CD107a+ CD8^+^ T_CM_ cells in the PCS vaccine group demonstrated that immunization with the PCS vaccine generated PCS specific CD8+ central memory T cells that have better potential in killing viral infected cells.

We conducted phenotypic analysis of Th17 and CD8+Th17 cells that were considered to be susceptible target cells for HIV or inflammatory disease^55-57^. Under *ex vivo* conditions, monkeys of the PCS vaccine group had a lower proportion of naive Th17 cells (CD4^+^IL-17A^+^CCR7^+^CD45RA^+^) and naive CD8^+^ IL-17A^+^ T cells expressing SIV/HIV co-receptor CCR5 (Figure 3G-H). They also had a lower proportion of naive CD8^+^IL-17A^+^ T cells expressing activation markers CD38 and CD69 (Figure 3I, 3J). With PCS peptide stimulation the monkeys in the PCS vaccine group also had a lower proportion of PCS peptide specific Th17_EMRA_ and CD8^+^IL-17A^+^ T_CM_ cells expressed CD107a (Figure 3H, 3I). Thus, immunization with the PCS vaccine generated fewer viral susceptible target cells and fewer CD8^+^IL-17A^+^ T cells that were involved in inflammatory disease^55-57^.

Antigen-specific regulatory T cells protect against aberrant immune responses^51^. We analyzed vaccine induced T regulatory cells. The monkeys in the PCS vaccine group had a higher frequency of PCS peptide specific CD4^+^CD25^+^FOXP3^+^CD45RA^+^CD127^−^ (Treg _EMRA_) cells and a higher frequency of PCS peptide specific CD4^+^CD25^+^FOXP3^+^CD45RA^+^CD127^+^ (naive Treg) cells expressing immunoregulatory cytokine IL-10 compared to that of the control group (Figure 4J, 4K). Taken together, the monkeys of the PCS vaccine group had lower immune activation, lower activated PCS peptide specific CD4^+^ T cells – the primary SIV target cells, but higher PCS peptide specific CD8^+^ T memory and CD4^+^ Treg responses.

### Immunologic correlates of protection (CoPs) induced by vaccination

We conducted regression analysis to identify potential immunologic factors that might be positively correlated with protection against acquisition of SIVmac251 infection in the vaccinated group. We assessed immunological measurements, including CVL mucosal inflammatory cytokines, CVL and plasma antibodies, PBMC cytokine-producing antigen recall responses and CD4^+^ or CD8^+^ T cell subsets (memory, IL-17A^+^ and Treg, *ex vivo* or Ag recall) from the peak (week 73) and pre-challenge (week 90) time points. The Spearman rank correlation was first used to assess individual immunological correlate of protection (CoP) observed in the PCS vaccine group, defined as the number of challenges required for SIVmac251 infection. The identified CoPs (Table 1) (positive Spearman’s rho with p<0.05, unadjusted for multiple inference) include (1) cytokines secreted by PBMCs (peak time point) after PCS peptide stimulations, including RANTES and IL-6 (to PCS1-4 peptides), and MIP-1α (to PCS9-12 peptides); (2) the frequency of CCR5 expressing Th17 T_EM_ (CD4^+^IL17A^+^CCR7^−^CD45RA^−^) cells (pre-challenge time point, *ex vivo*); (3) the frequency of CCR5 expressing CD8^+^IL17A^+^ T_EM_ cells (CD8^+^IL17A^+^CCR7^+^CD45RA^−^) and the intensity of CCR5 expression on these cells (pre-challenge time point, *ex vivo*); and (4) the frequency of PD-1 expressing PCS peptide specific Treg effector memory cell population (CD4^+^CD25^+^FoxP3^+^CD45RA^+^CD127^−^) (pre-challenge time point). These results suggested that protection was significantly correlated with T-cell responses. However, no significant associations with antibody response were observed.

**Table 1.** Immune correlates of Protection (CoPs)

| Immune response |  |  |  | Correlation with No. of challenges (Spearman rank) |  |
| --- | --- | --- | --- | --- | --- |
| Type | Sub-type | Ag | Time point | Rho (r) | p |
| Cytokine secretion (Ag recall) | RANTES | PCS1-4 peptide pool | Peak | 0.8500 | 0.0075 |
| Cytokine secretion (Ag recall) | MIP-1α | PCS9-12 peptide pool | Peak | 0.7798 | 0.0225 |
| Cytokine secretion (Ag recall) | IL-6 | PCS1-4 peptide pool | Peak | 0.7148 | 0.0463 |
| CD4+IL17+ T cells (ex vivo) | CCR5+% of CD4+IL17+CCR7-CD45RA- | N/A | Pre-challenge | 0.7750 | 0.0239 |
| CD8+IL17+ T cells (ex vivo) | CCR5+% of CD8+IL17+CCR7+CD45RA- | N/A | Pre-challenge | 0.8539 | 0.0070 |
| CD8+IL17+ T cells (ex vivo) | CCR5 MFI of CD8+IL17+CCR7+CD45RA- | N/A | Pre-challenge | 0.7563 | 0.0299 |
| CD4+CD25+FoxP3+ T cells (Ag recall) | PD-1+% of CD4+CD25+FoxP3+CD45RA+CD127- | PCS1-12 peptide pool | Pre-challenge | 0.8660 | 0.0054 |

Immune factors significantly correlating with number of challenges needed to cause infection were determined by Spearman rank correlation analysis. Ag: antigen (used to quantify antibody or to stimulate cells). CVL: cervicovaginal lavage wash. N/A: not applicable. Peak time point: Week 73 of vaccination schedule (1 week after final vaccination boost). Pre-challenge time point: Week 90 of vaccination schedule. Ab: antibody. Ex vivo: peripheral blood mononuclear cells (PBMCs) were not stimulated with Ag. Ag recall: PBMCs were stimulated with Ag in culture. MFI: mean fluorescent intensity. All analyses were conducted in the PCS vaccine group (n=8).

Several immune correlates of risk (CoRs) (defined as negative Spearman’s rho, with p<0.05 unadjusted for multiple inference) were also identified, including CVL mucosal IL-1α level at the peak and pre-challenge time points, CVL mucosal antibodies to PCS1, 3, 4, 5 and 9 at the peak time point, as well as the frequency or intensity of several activation or differentiation makers on subsets of CD4^+^ or CD8^+^ memory, IL17A^+^ or Treg cells at the pre-challenge time point (Table S7). These CoRs appear to relate to inflammation, immune activation and SIV target cell availability.

To explore more potential CoP/CoRs, especially combinations of immunological predictors which might act together to protect the vaccinated animals, we performed a multiple regression analysis of the immunological data using the LASSO (least absolute shrinkage and selection operator) (Figure 5). Although the risk of overfitting is large, and R^2^ optimistic (R^2^ = 0.964) in this situation, the LASSO regression model has nevertheless identified 6 potential predictors from pre-challenge T cell subsets, 5 of which were not identified via the single predictor Spearman correlation analysis (Figure 5A, 5B and Table S8). Individually these predictors also positively correlated with many other immunological variants by Spearman rank correlation analysis (Figure 5C and Table S8). The three predictors with positive coefficients in the LASSO model were positively correlated with several cytokine-producing antigen recall responses at the peak or pre-challenge time point. They were also positively correlated with the frequency or expression intensity of CD107a, TNF-α, MIP-1β, CD38, IFN-γ, IL-2 or CD69 in several T cell subsets (Th17_EM_, Th17_CM_, CD8^+^IL-17A^+^ T_CM/TEM_, CD8^+^ T naive/TEM, CD4^+^ T_EMRA_, CD4^+^/CD8^+^ Treg naive or CD4^+^Treg RM) either under *ex vivo* conditions or after PCS peptide stimulations at the pre-challenge time point (Fig. 5C and Figs. S7-9). The predictor CD4TregEmPD1, strongly correlated with 14 variants. Seven of the 14 variants plus the predictor itself also significantly correlated with vaccine protection, individually. The three predictors with negative coefficients in the LASSO model positively correlated with the fold change of two CVL mucosal cytokines, RANTES and IL-17A, at the pre-challenge time point, and with CVL antibodies to PCS4 and PCS5 at the peak time point (Figure 5C and Figs. S10-12). The predictor CD4TreNPD1m appeared to play an important role in regulating proliferation of several T cell subsets, as shown by its significant correlation with Ki67 frequency or/and expression intensity in CD4^+^ and CD8^+^ Tnaive, T_EM_, T_CM_, and T_EMRA_ cells under *ex vivo* conditions or after PCS peptide stimulations. CD4TreNPD1m also significantly correlated with the frequency or/and expression intensity of CD38, IFNγ, PD-1 in CD8^+^IL17A^+^ Tnaive, CD8^+^ T_EMRA_, and CD4^+^ Treg EM under *ex vivo* conditions and/or after PCS peptide stimulations. The other two predictors with negative coefficients in the LASSO model, CD8TregCmIL10 and CD4TreCmCD38 significantly correlated with the frequencies of CCR5^+^ Th17 naive cells after PCS peptide stimulation. The frequencies of these two central memory Treg cells significantly correlated with the frequency or expression intensity of IL-10, CD38, CD69, MIP-1β, and PD-1 in CD4^+^ Treg naive, CD4^+^ Treg EM, CD8^+^Treg naive, CD8^+^ Treg RM, CD8^+^ Treg EM, or CD8^+^ Treg CM cells under *ex vivo* conditions or after PCS peptide stimulations. These two features also correlated with the expression intensity of CD107a in CD8^+^IL17A^+^ T naïve, CD8^+^IL17A^+^ T_EM_ and Th17 T_EMRA_ cells, as well as the expression intensity of CD38 and IFNγ in Th17 T_EMRA_ cells.

**Figure 5.**
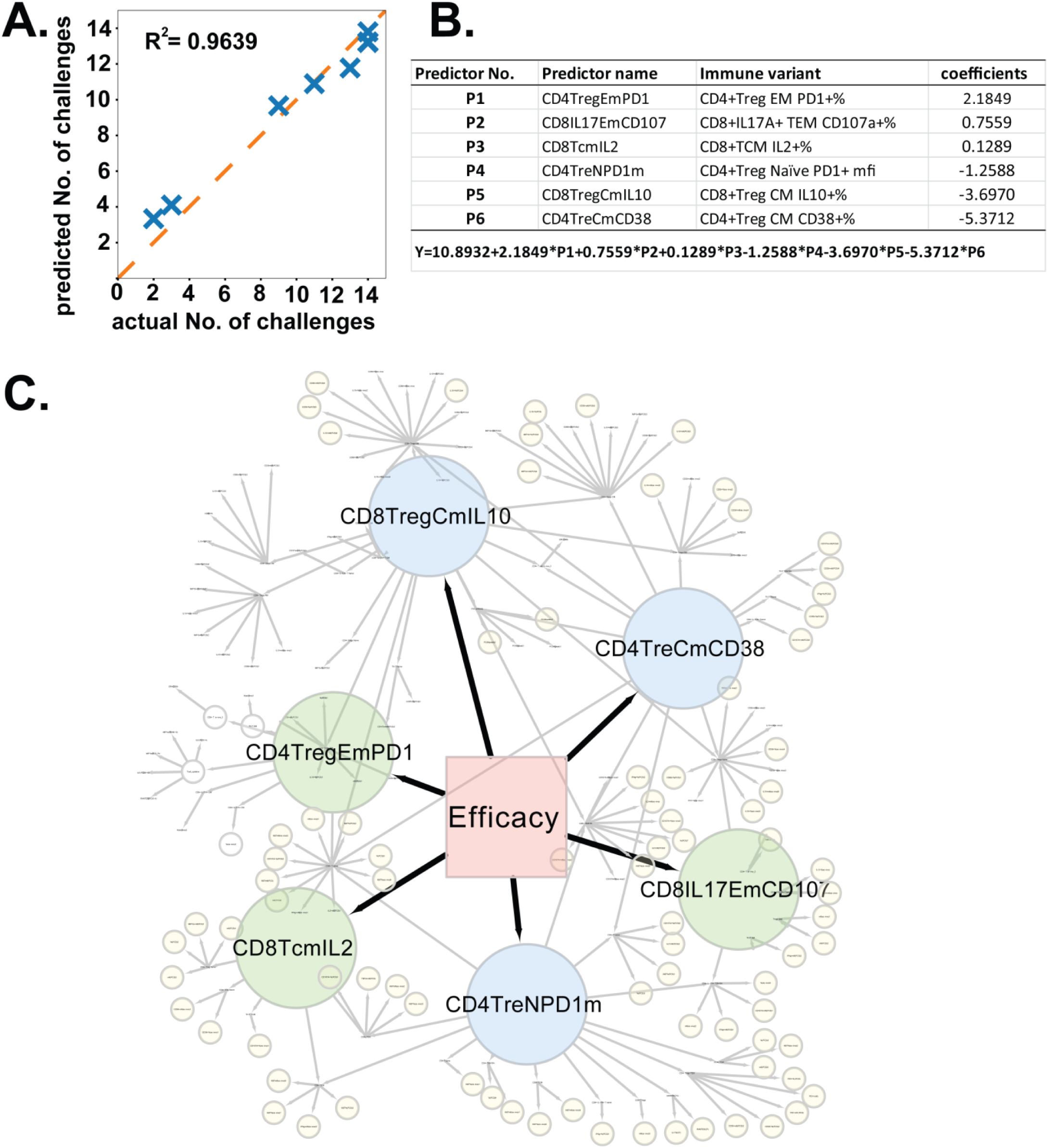
Immune predictors identified by LASSO analysis predict vaccine efficacy. a. LASSO model: Plot of actual versus predicted numbers of challenges from the identified immune predictors; b. The names of identified predictors and their coefficients by LASSO analysis. +/- signs indicate positive or negative association with protection, respectively. c. Networks of the identified immune predictors with other immune variants positively correlated the identified predictors by Spearman rank analysis (Table S8). The 6 identified predictors are labeled as big circled nodes in different colors. Green: predictors with positive co-efficient; Blue: predictors with negative co-efficient. Close-up view of the networks is available through the online version of this figure and in the Figs. S7-12.

Of the six predictors identified by the LASSO model, four are PCS peptide-stimulated Treg cells. These antigen-specific Treg cells appear to act together with PCS peptide specific IL2+CD8+ T central memory cells and PCS peptide specific CD107a+CD8^+^IL-17^+^ T effector memory cells to regulate the immunological environment in the vaccinated macaques and contribute to the effective protection against pathogenic SIVmac251 acquisition.

## Discussion

The development of an effective HIV vaccine is faced with great challenges. HIV has the formidable capacity to mutate its genome and evolves rapidly to evade immune recognition^58-60^. It targets CD4^+^ T cells and thrives in an activated immune system, a unique character distinguishing it from many other infectious pathogens^4,5^. The knowledge gained from the past failures and modest success in HIV vaccine development^61^ and from the natural immunity of highly exposed seronegative individuals^8^ are important in developing an effective HIV vaccine. Traditional candidate vaccines have been based on full HIV proteins such as Env and Gag, which contain multiple immunodominant decoy epitopes that divert immune responses away from targeting protective epitopes^62,63^. Furthermore, most of these vaccines may not have considered inflammation and immune activation associated with the strong and broad immune responses aimed by these vaccines. The potential impact of which may be an increase in activation of CD4 T cells, the preferred HIV targets, thus enhancing viral propagation and diminishing the benefit of anti-HIV immune responses. It has been proposed that the increased risk of infection observed in the vaccinees of the Step (HVTN 502) and Phambili (HVTN 503) trials, may have resulted from increased activation of mucosal CD4 T cell following vaccination^6,64,65^. The hypothesis is supported by preclinical data of similar vaccine studies in nonhuman primates (NHP)^6,64,65^. In the RV144 trial, the ALVAC vaccine used for prime was found to have much lower immunogenicity than the highly immunogenic Ad5 vaccine in the Step trial^66,67^. An NHP study mimicking the RV144 trial and its follow-up African trial observed that replacing the vaccine adjuvant alum with more immunostimulatory MF59 resulted in loss of the protective effect^68^. These data need to be considered in development and evaluation of anti-HIV vaccines.

Our study showed that the PCS vaccine addressed these concerns and provided greater than 80% protection of Mauritian cynomolgus macaques (MCM) against each of the repeatedly intravaginal SIVmac251 challenges. Cynomolgus macaques and Mauritian Cynomolgus macaques are susceptible to SIV and SHIV infection, its natural course of infection leading to simian AIDS more closely recapitulates the HIV infection in humans, and is a valid vaccine evaluating model ^22,24,69-81^. A study showed that there was no difference in terms of infectivity and peak viral load levels between rhesus macaques and Cynomolgus macaques inoculated through oral and rectal mucosal routes^82^.

Furthermore, the SIVmac251 is a swarm of viruses with high pathogenicity and significant genetic heterogeneity^81^. We believe that the MCM/SIVmac251 vaccine challenge model is stringent in evaluating the vaccine efficacy. The efficacy of the PCS vaccine is among the best protective vaccines thus far tested in the NHP models^23,27,83^. The PCS vaccine focused immune responses on the functionally critical part of the virus – sequences surrounding the protease cleavage sites. Anti-PCS peptide CD8^+^ central memory T cells with higher potential of killing viral infected cells were among the T cell responses correlated with vaccine efficacy. The mucosal inflammation induced by the PCS vaccine is low. The amount and activation of viral target T cells in the blood was significantly lower in the PCS vaccine group than that in the control group. The PCS vaccine immunized monkeys had significantly lower CCR5 expression in Th17 cells, lower frequencies of CCR5^+^IL-17A^+^CD8^+^ T naive cells, and lower expression of activation markers CD69 and CD38. These are relevant to HIV infection since Th17 cells are preferential targets for productive infection^55,56^ and IL17A^+^CD8^+^ T cells also are thought to play a role in immune pathology that contributes to the maintenance of target cell populations for HIV infection^57^. Thus, the PCS vaccine limited the availability of viral target cells. Furthermore, the PCS-vaccinated monkeys had higher frequency of PCS-specific CD4^+^ Treg EMRA and CD4^+^IL-10^+^ Treg naïve cells than controls, thus they may be better able to regulate anti-viral immune responses. Six predictors identified using the LASSO model that predicted vaccine efficacy are all PCS peptide specific T cell responses including four PCS peptide-specific Treg cell populations and two PCS peptide specific CD8 T central/effector memory cell populations. It suggests that regulation of anti-SIV CD8 T cell responses is important.

The PCS vaccine generated a focused and modest magnitude of protective immune responses. These antiviral immune responses in combination with low inflammation and reduced CD4^+^CCR5^+^ viral target cells appear to be sufficient to prevent pathogenic SIVmac251 acquisition. Since most sexual HIV transmission was initiated by a single founder virus^84,85^, a focused, modest-magnitude and well-regulated antiviral immune response might be enough to prevent the viral establishment and propagation as demonstrated in this study.

In conclusion, this study showed that a novel HIV vaccine candidate focusing immune response to sequences surrounding the 12 protease cleavage sites can protect female macaques from pathogenic SIVmac251 intravaginal acquisition. For the first time, a candidate HIV vaccine without full Gag and Env proteins as immunogens has demonstrated high efficacy of protection. Our study also showed that the PCS immunogen-specific CD8^+^ T central and effector memory cells, and PCS immunogen-specific CD4^+^ and CD8^+^ regulatory T cells correlated with the vaccine efficacy. Thus, an effective prophylactic HIV vaccine needs to not only generate effective antiviral immune responses to kill infected cells, but also regulate the immune responses and control immune activation and inflammation. These can be achieved by the PCS vaccine, a vaccine targeting HIV maturation.

In order to observe within a reasonable time that the vaccine was effective, the repetitive low-dose SIVmac251 intravaginal challenges using the NHP model employed virus titres far exceeding the average HIV viral load in semen^86-88^. This ensured that the NHP/SIVmac251 model used in this study be highly stringent. Although this study shows that the PCS vaccine is effective in preventing infection in the setting of a highly stringent NHP/SIVmac251 intravaginal challenge model, human clinical trials will constitute the ultimate test. The detailed available information of T cell epitopes surrounding the 12 HIV protease cleavage sites of major HLA class I alleles in the world populations^11^, and the immune correlates of protection identified in this study will facilitate future clinical trials.

## Materials and Methods

### Study design

Vaccine efficacy against acquisition of infection was evaluated in female MCMs receiving repeated low-dose (RLD) intravaginal challenges by SIVmac251, which mimics natural HIV infection through the female genital tract. Sample sizes were determined using the parameters described in previous studies^89^. An infection rate of 0.4 among control macaques and an interval of two weeks among each challenge of mucosal RLD were estimated from our pilot study. Statistical power was then calculated based on assuming an infection rate of 40% per exposure and a susceptibility to infection of 85% in control animals at the end-point of the 6^th^ viral challenge when majority of control macaques become infected. To achieve 80% statistical power with a 2-sided type I error rate < 0.05, a total of 16 female MCMs were required and assigned into two groups (eight per group): the PCS and sham control vaccine groups.

### Experimental animals and ethics statement

Female Mauritian cynomolgus macaques (MCMs) were pair-housed within the same experimental group during the immunization phase of the study with visual and auditory access to other conspecifics. Paired monkeys lived in two adjacent standard stainless-steel primate cages (27”L × 27”W × 32”H per cage). Rooms were maintained at 65–75°F, 30–70% humidity, and on a 12:12 light-dark cycle (ON: 0600, OFF: 1800). Standard nonhuman primate chow with fruit or vegetables was provided daily. In addition, we provided foraging activities and physical environmental enrichment at least weekly for both activities. All animals were observed at least twice daily for health or welfare issues. Sedation (ketamine alone, or ketamine/dexmedetomidine, atipamezole for reversal) was provided during the experimental procedures. The experiments were approved by the University of Wisconsin IACUC protocol (G005765) in accordance with the US Animal Welfare Act and following the recommendations of the National Research Council Guide for the Care and Use of Laboratory Animals, 8th Edition and the Weatherall report, The Use of Nonhuman Primates in Research. The Wisconsin National Primate Research Center is fully accredited by AAALAC under the University of Wisconsin, Division of Vice-Chancellor for Research and Graduate Education.

All animal work was carried out at Wisconsin National Primate Research Center. We conducted vaccine evaluation using the Mauritian cynomolgus macaque (MCM)/SIVmac251 infection model. MCMs are abundant in supply, and have simple major histocompatibility complex (MHC) genotypes (with only seven MHC haplotypes^22,69,90-94^). Their infection by SIVmac251 leads to peak and set point viral loads and disease progression pattern that closely mimic human HIV infection^22,95,96^. This infection model has proven stringent for testing prophylactic vaccines: None of the vaccines previously tested in this model has demonstrated significant efficacy against acquisition of SIVmac251 (or SIVmac239) infection^22,24,72-76,78-80,97-107^. Additional animal information, concerning grouping, ID, gender, age, weight and MHC haplotypes, is summarized (Table S1-S2).

### Immunization scheme

The PCS vaccine contained twelve 20-mer peptides, derived from the twelve protease cleavage sites (PCSs) of SIVmac239 (ten amino acids flanking each side of the cleavage site)^12,108^. These peptide immunogens were delivered as recombinant vesicular stomatitis viruses (rVSVpcs) or biodegradable nanoparticles (NANOpcs)^12^. The rVSVpcs consisted of twelve different rVSV viruses each delivering one PCS peptide (PCS1, PCS2, …, or PCS12) and the NANOpcs was formed by a pool of twelve formulations, each made of chitosan/dextran sulfate nanoparticles containing one different PCS peptide^12^. The rVSVpcs and NANOpcs in combination were referred to as the PCS vaccine. The sham control vaccine was rVSV vector (rVSVvt) virus (without SIV immunogen insert) and sterile water (NANO vehicle)^12^. The vaccination scheme was described previously^12^, consisting of a prime (with rVSV at week 0) and four boosts (1^st^ boost: rVSV + NANO at week 6, 2^nd^ boost: NANO at week 16, 3^rd^ boost: rVSV + NANO at week 51, and 4^th^ boost: rVSV at week 72). rVSVs were administered intramuscularly at a dose of 2×10^7^ pfu/animal (i.e. 1.67×10^6^ pfu/rVSVpcs type/animal × 12 PCS types = 2×10^7^ pfu/animal for the PCS vaccine group, or 2×10^7^ pfu of rVSVvt/animal for the control group), except that, in the 4^th^ boost, 1×10^8^ pfu of each rVSVpcs or rVSVvt was used. NANOpcs was administered intranasally at a dose of 50 µg peptide/PCS type/animal × 12 PCS types = 600 µg/animal.

### Repetitive low-dose intravaginal SIVmac251 challenges

Challenges were carried out every two weeks starting 24 weeks after the last vaccination boost. The vast majority of SIV challenges in previous studies were performed weekly^23,26,109^ while viral acquisition status was monitored by quantification of plasma viral load (VL). However, establishment of systemic infection can take up to 10 – 14 days^54^. Potentially some slowly establishing infections may not be detectable within one week post challenge by the routine method of plasma VL measurement. This was also confirmed in our study. Therefore, using the weekly challenge protocol, an infection identified during a later challenge round may actually have been acquired in an earlier round, leading to an over-estimation of vaccine efficacy. In this regard, we carried out challenges every two weeks, and plasma VL was quantified on days 6, 10 and 14 of each challenge round (any animal found infected was not challenged further). Indeed, we found that in some cases an infection could remain below detectable level until as late as 14 days following a challenge (Table S3). In addition, we took into consideration that in females, susceptibility to vaginal HIV infection could be affected by phases of the menstrual cycle. It was reported that the susceptibility is higher in the luteal phase than in non-luteal phases^110-114^. Our 2-week challenge cycle was also designed to maximize the chance for even distribution of challenges in the luteal versus non-luteal phases, since the luteal phase constitutes approximately half of a 4-week menstrual cycle. It was confirmed that in both the control and PCS vaccine groups challenges fell in the luteal and non-luteal phases nearly equally, without any significant difference between the two groups (Figs. S1 and S2).

Intravaginal challenges were performed by atraumatically delivering 250 times 50% tissue culture infectious doses (TCID50) of SIVmac251 in 1 ml of saline or tissue culture medium to the vagina while the animal was anesthetized as described below. The virus was delivered using a TB syringe with the needle removed (or using a TB syringe with a rounded end gavage tube). The syringe was gently inserted into the vagina about 4cm, then withdrawn slightly and the dose was injected slowly over 1 minute, then the animal’s pelvic region kept elevated up to 30 minutes before returning the animal to its cage. When withdrawn, the syringe was examined to ensure that there was no trauma to the vagina inflicted by the syringe (in which case blood would be visible).

For anesthesia, animals were immobilized against the front of the cage by a squeeze back mechanism, and anesthetized using up to 7 mg/kg ketamine (i.m.) and up to 0.03 mg/kg dexmedetomidine (i.m.) to be reversed at the conclusion of a procedure by up to 0.3 mg/kg atipamezole (i.v. or i.m.). Any additional anesthesia was administered only in consultation with a WNPRC veterinarian. Alternative anesthesia would only be used as directed by a WNPRC veterinarian. The duration of anesthesia was usually less than 45 minutes. Monitoring of anesthesia recovery was documented every 15 min until the animal was sitting upright, then every 30 minutes until the animal was fully recovered from the anesthesia.

### Viral load analysis

Viral RNA was isolated from plasma samples using the Maxwell 16 Viral Total Nucleic Acid Purification kit on the Maxwell 16MDx instrument (Promega, Madison WI). Viral RNA was then quantified using a highly sensitive qRT-PCR assay based on a previously published protocol^115^.

### Bio-Plex multiplexed cytokine/chemokine assay

We used a customized cytokine/chemokine panel targeting 14 (pro- or anti-) inflammatory cytokines/chemokines including Granulocyte Macrophage Colony Stimulating Factor (GM-CSF), Macrophage Chemo-attractant Protein 1 (MCP-1), Monocyte Inflammatory Protein 1 alpha (MIP-1α), Monocyte Inflammatory Protein 1 beta (MIP-1β), CCL5/RANTES, IFN-γ inducible protein 10 (IP-10/CXCL-10), Tumor Necrosis Factor alpha (TNF-α), Interferon gamma (IFN-γ), IL-1α, IL-8, IL-10, IL-17a, IL-1β, and IL-6. Mucosal inflammatory responses and cellular recall responses (see below) were measured using cervicovaginal lavage (CVL) and PBMC culture supernatant samples, respectively. Antibodies and protein standards used in these assays are listed in Table S9. The CVL sample collection method was described previously^12^. For the CVL assay, PBS with 1% BSA (Assay buffer) was used both as general diluent and blank controls, while RPMI-1640 Culture Medium (Sigma, R8758-6X500ML) was used as diluent for standards and blank controls for the PBMC supernatant samples. In order to generate the standard curves, we prepared a cytokine/chemokine protein standard pool with final concentrations listed in Table S10, using PBS with 1% BSA as diluent. We then prepared eight four-fold serial dilutions of this cytokine/chemokine protein standard pool and transferred 50 µL of each dilution to a 96-well Bio-Plex Pro™ Flat bottom plate (BioRad, Catalog# 171025001). Final standard curves were optimized and generated using Bio-Plex Manager 6.1 software, where coefficient of variation (%CV) was no more than 30% and percentage of recovery was at least 70. 20 µg of capture antibody for each cytokine/chemokine type was coupled to 1.25×10^7^ Bio-Plex Pro™ Magnetic COOH Beads (BioRad, Catalog# MC10053-01) using Bio-Plex Amine Coupling Kit (Bio-Rad, Catalog# 171-406001) according to the manufacturer’s suggestions and diluted to a final concentration of 5×10^6^ beads/mL. 50 µL of undiluted sample was added to each well containing ∼420,000 coupled magnetic beads/capture antibody type (1 in 600 dilutions) and incubated while shaking at high speed (∼1000rpm) for 30 seconds, then at medium speed (∼300rpm) for one hour on a plate shaker at room temperature. After washing the plate 3× (MAG3× setting on BioRad wash station) with wash buffer (PBS with 0.05% Tween20), 30 µl of biotinylated detection antibody at 1µg/ml was added to each well and again incubated while shaking for 30 minutes at room temperature. After another routine wash, 50 µl of Streptavidin-PE in 1:100 dilution of BioRad 1× stock (Catalog# 171304501) was added per well and incubated while shaking for 10 min at room temperature. A final wash was performed and afterwards 125 µl of assay buffer was added to each well. After shaking the plate for 10 seconds, we ran the plate on a Bio-Rad Bio-Plex™ 200 System. Each bead fluorescence intensity and corresponding protein concentration (pg/ml) were generated by Bio-Plex Manager 6.1 software. To take into account dilution variations during CVL sample collections, cytokine/chemokine concentrations were all normalized to total protein concentrations in the same sample. CVL total protein concentration was quantified using NanoOrange® protein quantitation kit (Thermo Fisher Scientific) according to the supplier’s protocol.

### MHC typing and Bio-Plex multiplexed antibody assay

These were described previously^12,108^.

### IFN-γ enzyme-linked immunospot (ELISPOT) assay

Freshly isolated PBMCs (2-2.5 × 10^5^/well) were stimulated for 16-18 hours in a 96-well pre-coated plate (ELISpotPLUS kit, MABTECH Inc., Mariemont, OH) at 37°C in a 5% CO_2_ incubator. Stimulation was performed with PCS peptide pool, targeting subsets of PCSs (PCS1-4, PCS5-8 and PCS9-12, respectively), at 2.5μg/ml per individual peptide. The sequences of these peptides (Sigma-Aldrich) were listed in Table S11. Cells stimulated with 5.0 µg/ml of concanavalin A (Sigma Chemical, St. Louis, MO) were used as positive control. All tests were performed in duplicate. Wells were imaged with an AID ELISPOT reader (AID, Strassberg, Germany), and counted by AID EliSpot Reader version 3.2.3, with set parameters for spot size, intensity and gradient. Background (mean of wells without peptide) levels were subtracted from each well on the plate. A response was considered positive if the mean number of SFCs of duplicate sample wells exceeded twice the background and was >50 SFC per 1 × 10^6^ cells.

### Cell culture of PBMCs and PCS peptide stimulation for analysis of multiple cytokine secretions or T cell subsets

2.0×10^6^ PBMCs/ml in R10 media (RPMI1640-HyClone + 10% FBS + 2% antibiotic/antimycotic, Thermo Fisher Scientific) were added into each well of a 96-well plate, and cultured in media only (no-stimulation negative control), in the presence of 0.001µg/ml PMA with 0.01µg/ml Ionomycin (positive control), or with 1.5µM SIV PCS peptide pools (Sigma-Aldrich, Table S11), overnight at 37°C with 5% CO_2_. Cytokines secreted into the culture supernatants were analyzed by the Bio-Plex multiplexed cytokine assay as described above. For flow cytometry analysis (with intracellular staining) of T cell subsets, PBMC culture and stimulation were performed in the presence of protein transport inhibitors, 50nL GolgiStop™ (monensin) and 200nL GolgiPlug (brefeldin A).

### Flow Cytometry

PBMCs with minimal manipulation (ex vivo) or with stimulation by a total PCS peptide pool (PCS1-12, Table S11) were transferred to 12 × 75mm^3^ polystyrene tubes (BD Falcon) and washed with FACS wash (PBS supplemented with 2% FBS). Three sets of fluorochrome-labeled antibody cocktails (Table S9) were prepared, and added to the washed PBMCs, and incubated at room temperature for 30 minutes in the dark. The extracellular antibody cocktails panels included: Panel 1 – LIVE/DEAD™ fixable Blue stain (ThermoFisher Scientific), CD3 V500 SP34-2, CD4 BV605 L200, CD8 V450 RP8-T4, CCR7 A700 15053, CD127 PE-CF594 HIL-7R-M2 and CD45RA/RO APC-H7 UCHL1; Panel 2 (T cell regulatory and activation) – LIVE/DEAD™ fixable Blue stain, CD3 V500 SP34-2, CD4 BV605 L200, CD8 APC-Cy7 RP8-T4, CCR7 A700 15053, CD127 PE-CF594 HIL-7R-M2, CD38 FITC HB7, PD-1 BV650 M1H4, CD69 APC FN50, CD25 PE-Cy5 and CD45RA PE-Cy7 L48; and Panel 3 (Th17) – LIVE/DEAD™ fixable Blue stain, CD3 V500 SP34-2, CD4 BV605 L200, CD8 V450 RP8-T4, CD38 FITC HB7, CD69 APC FN50, CD45RA PE-Cy7 L48, CCR7 A700 15053, CD127 PE-CF594 HIL-7R-M2 and CD195 PE-Cy72 D7/CCR5 (all antibodies from BD Biosciences). Following the surface staining 500µl FACS wash was added to each tube to wash the cells followed by centrifugation at 1500rpm for 5 min with low brakes. Next, the cells stained with extracellular antibodies for panels 1 and 3 were permeabilized using 150µl Cytoperm/Cytofix™ (BD Biosciences) for 10-15 min, while those for panel 2 ICS staining were fixed using 2ml of 1x FoxP3 buffer (BD Pharmingen) for 10-15 min. The cells (panels 1-3) were then washed using FACS wash, and incubated with ICS panel 1 and 3 antibodies for 30 minutes. Alternatively, the cells (panel 2) were further permeabilized using 1X Human FoxP3 Buffer C (BD Pharmingen) for 30 min, then washed using FACS wash, and finally incubated with a cocktail of ICS antibodies. The panels of ICS antibodies used were as follows: Panel 1 ICS – CD107a PE-Cy5 H4:H3, IFN-γ BV786 4S:B3, IL-2 FITC MQ1-17H12, TNFα APC Mab11, Ki67 PE-Cy7 B56, MIP1α PE D21-1351 and IL-17A BV650 N49-653; Panel 2 – FoxP3 V450 259D/C7, IL-10 BV711 JES3-9D7 and MIP1α PE D21-1351. Panel 3 – CD107a PE-Cy5 H4:H3, IL-17A BV650 N49-653, IFN-γ BV786 4S:B3, and MIP1α PE D21-1351. Following the ICS antibody incubation, the cells (panels 1-3) were washed using FACS wash, and 300µl 1% formaldehyde with 1X PBS was added to each tube. Flow cytometry analysis was done on a multicolor BD LSRII cytometer with data acquisition using BD FACS DIVA software. Data analysis was carried out using FlowJo (courtesy of African HIV/AIDS program -Treestar).

### General statistical analysis

Time to infection in each group, measured by the number of challenges required to acquire infection and censored following challenge 6 on day 84, was summarized using the Kaplan-Meier survival curves and compared using the log-rank test; vaccine efficacy was evaluated as (1 – Hazard Ratio)*100% in a proportional hazard regression model. Spearman rank correlations were used to assess the associations between each immune factor and the number of challenges to infection. Immune responses were compared between the vaccine and control groups using Mann-Whitney’s U. For each of the above tests, p < 0.05 was considered significant, without adjustment for multiple inference.

### LASSO analysis for a predictive model of vaccine efficacy

Our Spearman rank analysis considered each immunologic measure individually. To identify potential combinations of immunological predictors of resistance or susceptibility, we also performed a multiple regression analysis. We used a linear model for the number of challenges survived versus the available biological measurements (features) due to the small numbers of monkeys in each group. The linear model was fitted using the LASSO, a variant of least-squares regression with an L1 regularization term, implemented in the Python scikit-learn library v0.20.3^116^. This regularization forces most of the coefficients in the final model to be zero and selects only the most important features, to a maximum of 7 features for a group of 8 animals, before estimating the final model. The regularization parameter was selected using a leave-one-out cross-validation strategy to achieve the lowest squared prediction error averaged across the fitted models. Each feature was scaled to range from 0 to 1 to facilitate the interpretation of the regression coefficients.

## Acknowledgments

We thank staff at Wisconsin National Primate Research Center Scientific Protocol Implementation Unit and Immunology Services Unit, especially Dr. Eva Rakasz for important technical support. Dr. Michael Seaman’s lab at BIDMC for plasma antibody neutralization assay. We thank Dr. Stuart Shapiro, NIH Vaccine Research Program, and Dr. Matthew Gilmour, National Microbiology Laboratory of Canada, for their support and discussion. We thank Dr. Jon Warren and Dr. Nancy Miller for providing SIVmac251 Desrosiers” 2010-Day 8 viral stock, and NIH AIDS Reagent Program for providing SIV and HIV antibodies, antigens and other important reagents. We thank Mr. Wallace Trenholm and Dr. Mark Alexiuk of Sightline Innovation, for supporting LASSO analysis. We express our appreciation to the support of Dr. Greg Hammond, Dr. Allan Ronald and Dr. Jo Kennelly for their support for this vaccine study.

## Authors contributions

M.L. designed and coordinated the study with N.S., M.J.A., Q.L., J.W., D.S., J.C., H.L., interpreted the data and wrote the manuscript. B.L., R.F.S., A.B.L., X.Q.L. analyzed the vaccine efficacy data, performed statistical analysis, LASSO analysis, and the correlates of protection/risk analyses, prepared the figures and helped to write the manuscript. R.W.O. performed the flow cytometry analysis for PBMCs in the blood, FACS data analysis, and helped to write the manuscript. H.L. prepared the rVSVs for vaccinations, performed the studies on antibodies together with L.L. and Y.H. and analyzed the antibody data and prepared some of the figures and tables, and drafted manuscript together with M.L.. N.T. performed the analysis of cytokines in the CVL and performed cytokine analysis with R.W.O. and M.K. and helped for data analysis. N.S. coordinated animal study with D.S. (who performed sampling, immunization, challenge experiments together with animal care technicians). M.J.A., J.C., J.F.P. and T.G.D. provided nanoformulated PCS vaccine for immunization and boost. Q.L. and Y.W. performed data acquisition and analysis. F.P., S.L., Q.L., J.W., and N.D. helped to write the manuscript. M.L., F.P., N.S., M.J.A. and J.W. secured funding for this study. L.L. helped for statistical analysis.

## Data and materials availability

The authors declare that the data supporting the findings of this study are available within the paper and its supplementary information files. The data that support the findings of this study are available from the corresponding author upon reasonable request. The materials can be made available upon request through an MTA.

