## Supplemental figures and tables for "A novel vaccine targeting the viral protease cleavage sites protects Mauritian cynomolgus macaques against vaginal SIVmac251 infection"

**This PDF file includes:**

Figs. S1 to S12
Tables S1 to S12

**Menstrual phase distributions of SIV challenges (standard)**

| Group | Animal ID | Number of challenges | | Time of infection |
|---|---|---|---|---|
| | | LT | N-LT | |
| Control | cy0774 | 1 | 1 | LT |
| Control | cy0775 | 1 | 1 | LT |
| Control | cy0776 | 0 | 1 | N-LT |
| Control | cy0777 | 2 | 2 | N-LT |
| Control | cy0778 | 3 | 3 | - |
| Control | cy0779 | 3 | 3 | - |
| Control | cy0780 | 2 | 2 | LT |
| Control | cy0781 | 2 | 0 | LT |
| PCS | cy0758 | 2 | 1 | LT |
| PCS | cy0759 | 3 | 3 | - |
| PCS | cy0760 | 3 | 3 | - |
| PCS | cy0761 | 3 | 3 | - |
| PCS | cy0762 | 3 | 3 | - |
| PCS | cy0763 | 3 | 3 | - |
| PCS | cy0764 | 1 | 1 | LT |
| PCS | cy0765 | 3 | 3 | - |

**LT: luteal phase; N-LT: non-luteal phase; - : not infected**

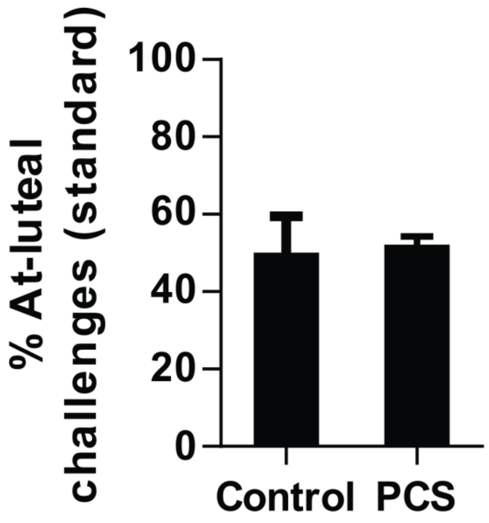


**Fig. S1 Menstrual phase distributions of SIV challenges.** Data were based on the standard challenge protocol (six challenges)

**Menstrual phase distributions of SIV challenges (extended)**

| Group | Animal ID | Number of challenges | | Time of infection |
|---|---|---|---|---|
| | | LT | N-LT | |
| Control | cy0774 | 1 | 1 | LT |
| Control | cy0775 | 1 | 1 | LT |
| Control | cy0776 | 0 | 1 | N-LT |
| Control | cy0777 | 2 | 2 | N-LT |
| Control | cy0778 | 7 | 6 | - |
| Control | cy0779 | 7 | 6 | - |
| Control | cy0780 | 2 | 2 | LT |
| Control | cy0781 | 2 | 0 | LT |
| PCS | cy0758 | 2 | 1 | LT |
| PCS | cy0759 | 7 | 6 | LT |
| PCS | cy0760 | 6 | 7 | - |
| PCS | cy0761 | 7 | 6 | - |
| PCS | cy0762 | 4 | 5 | LT |
| PCS | cy0763 | 6 | 7 | - |
| PCS | cy0764 | 1 | 1 | LT |
| PCS | cy0765 | 5 | 6 | LT |

LT: luteal phase; N-LT: non-luteal phase; - : not infected

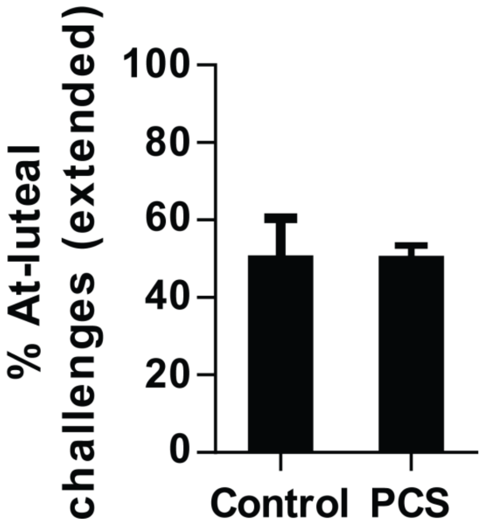


**Fig. S2 Menstrual phase distributions of SIV challenges.** Data were based on the extended (thirteen) challenges

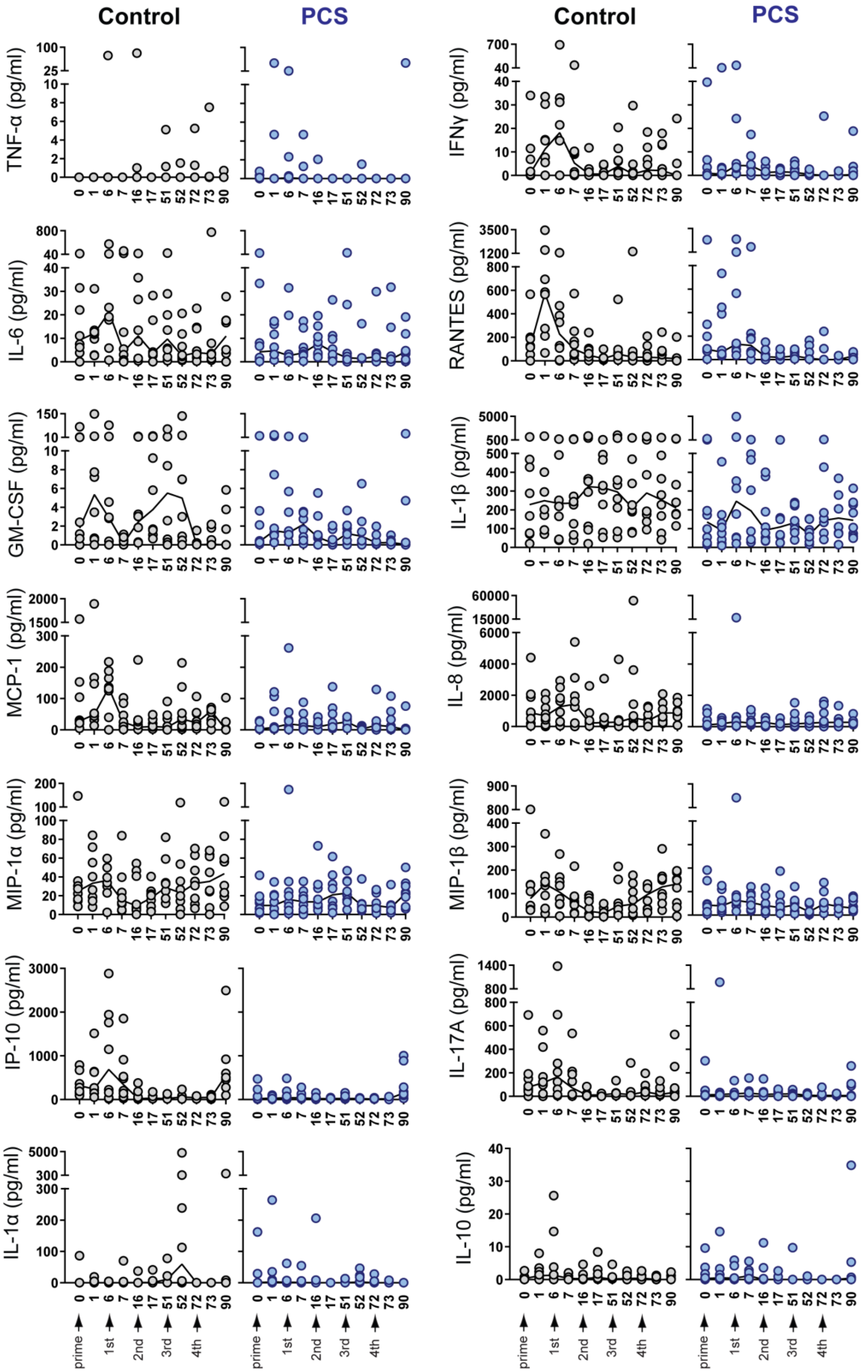


**Fig. S3 Dynamics of inflammatory cytokines in cervicovaginal secretions.** Cervicovaginal lavage (CVL) samples from the vaccination experiments illustrated in Fig. 1b were quantified by a Bio-Plex multiplexed cytokine assay, at indicated time points. Data are shown as values from individual animals with median line connection.

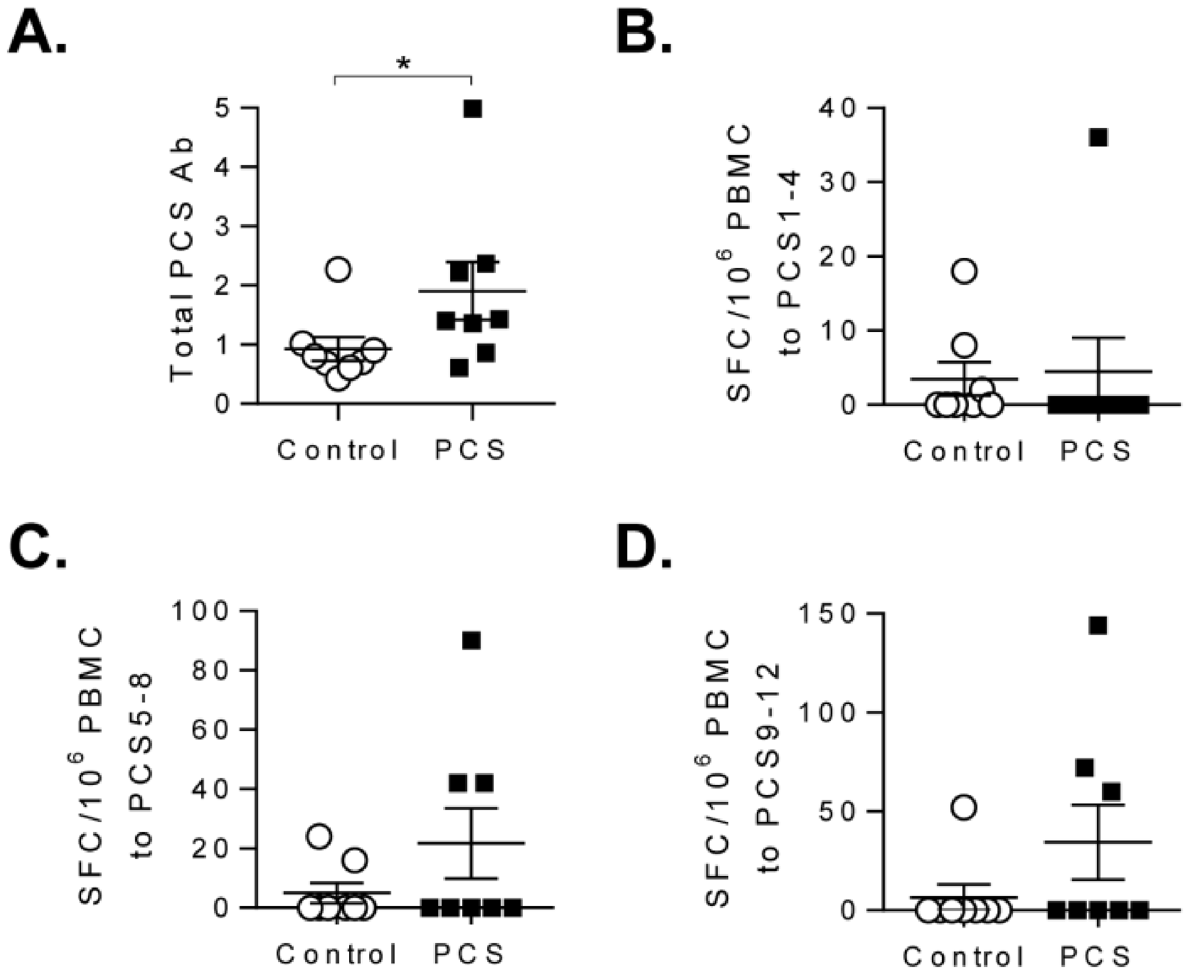


**Figure S4. The PCS vaccine induced anti-PCS antibody and IFN-γ ELISPOT responses.** (A). Plasma total anti-PCS antibodies at week 73 (peak time point). *$p < 0.05$ (Mann-Whitney's test). (B-D). Cellular antigen recall responses to PCS subset peptide pools in PBMCs isolated at week 73 were measured by IFN-γ ELISPOT assays. SFC, spot-forming cells. All data are presented as values from individual animals with mean ± SEM.

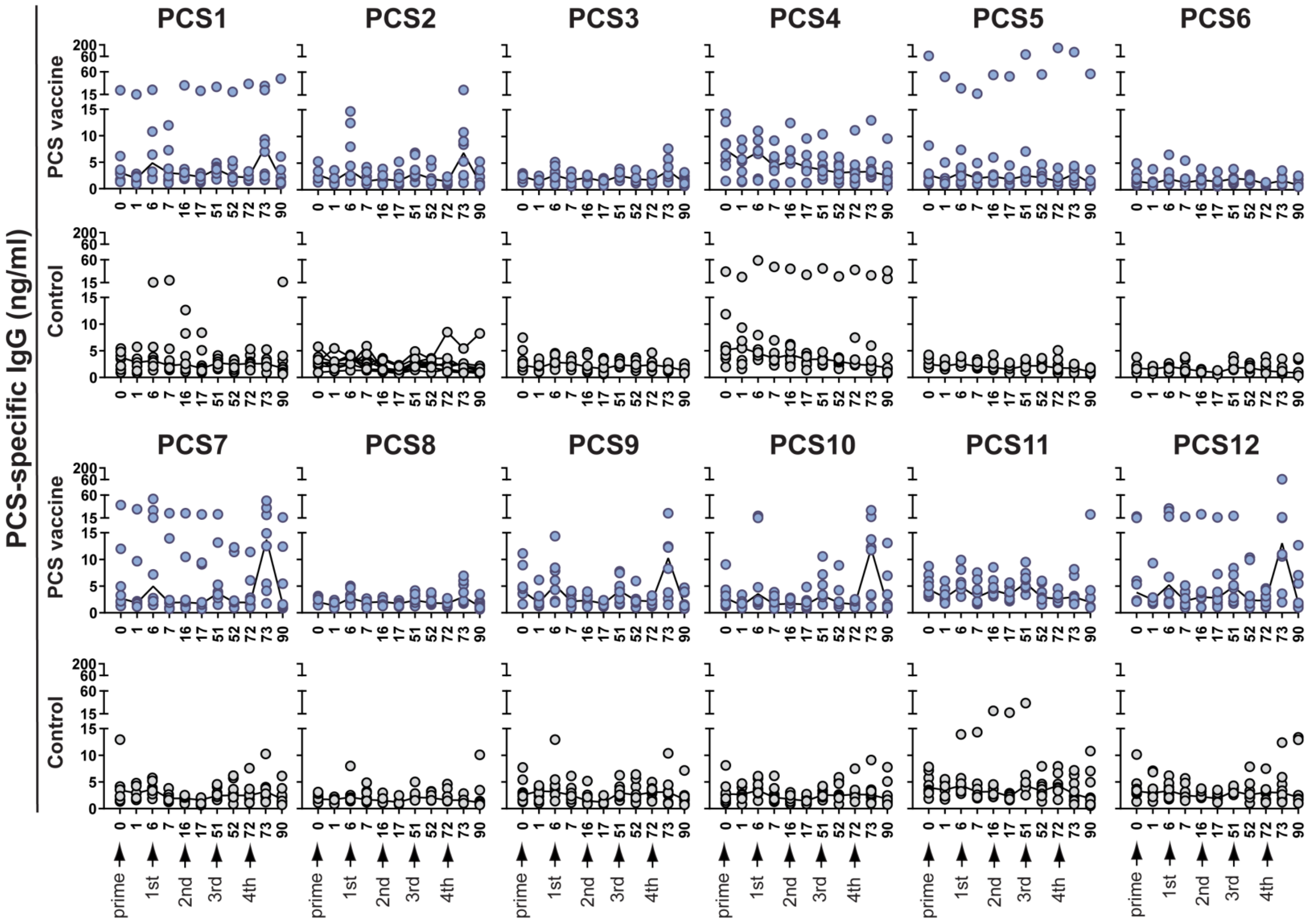


**Fig. S5 Dynamics of plasma IgG antibodies to PCS peptide antigens.** Plasma samples from the vaccination experiments illustrated in Fig. 1b were quantified for antibodies to each PCS peptide by a Bio-Plex multiplexed antibody assay, at indicated time points. Data are shown as values from individual animals with median line connection.

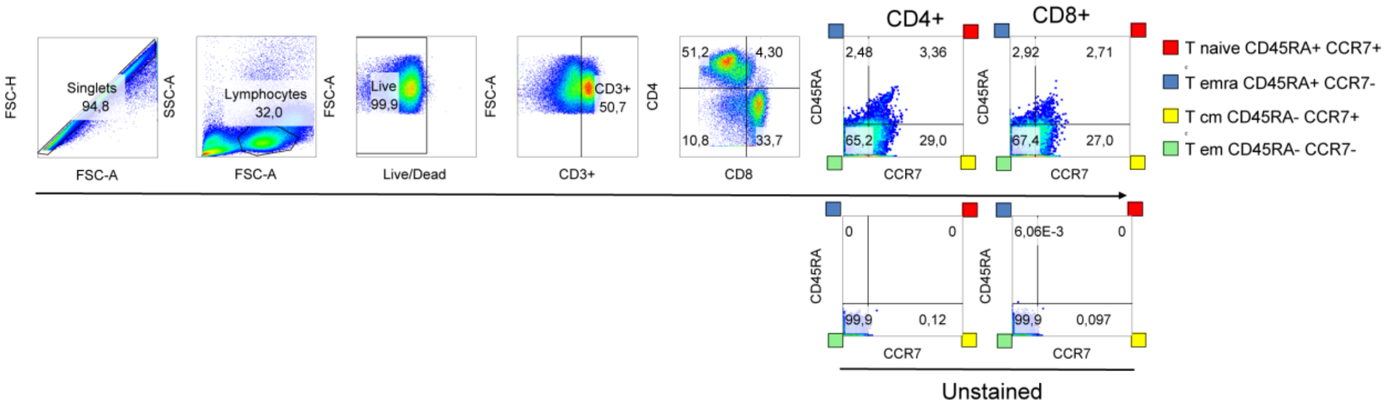


**Fig. S6** Representative of gating strategy in defining T cell subsets.

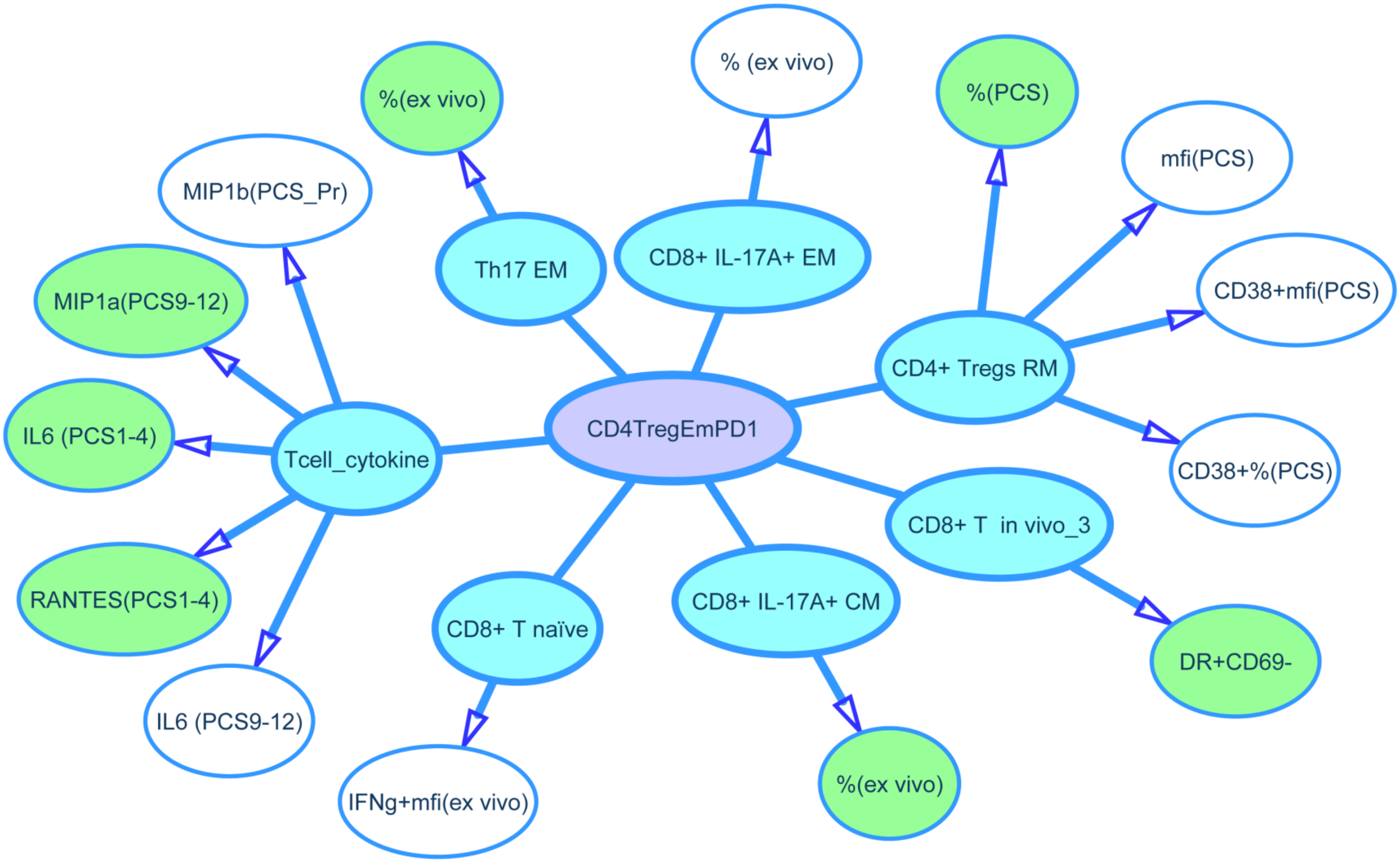


**Figure S7 Correlations of immune factors in relation to CD4TregEmPD1**. Different T cell subsets are in blue colored circles. Uncolored circles represent either frequency or expression intensity of immune activation or cytokine markers either under ex vivo conditions or after stimulated with PCS peptides. Green colored circles indicate variants that positively correlated with vaccine efficacy by Spearman rank analysis.

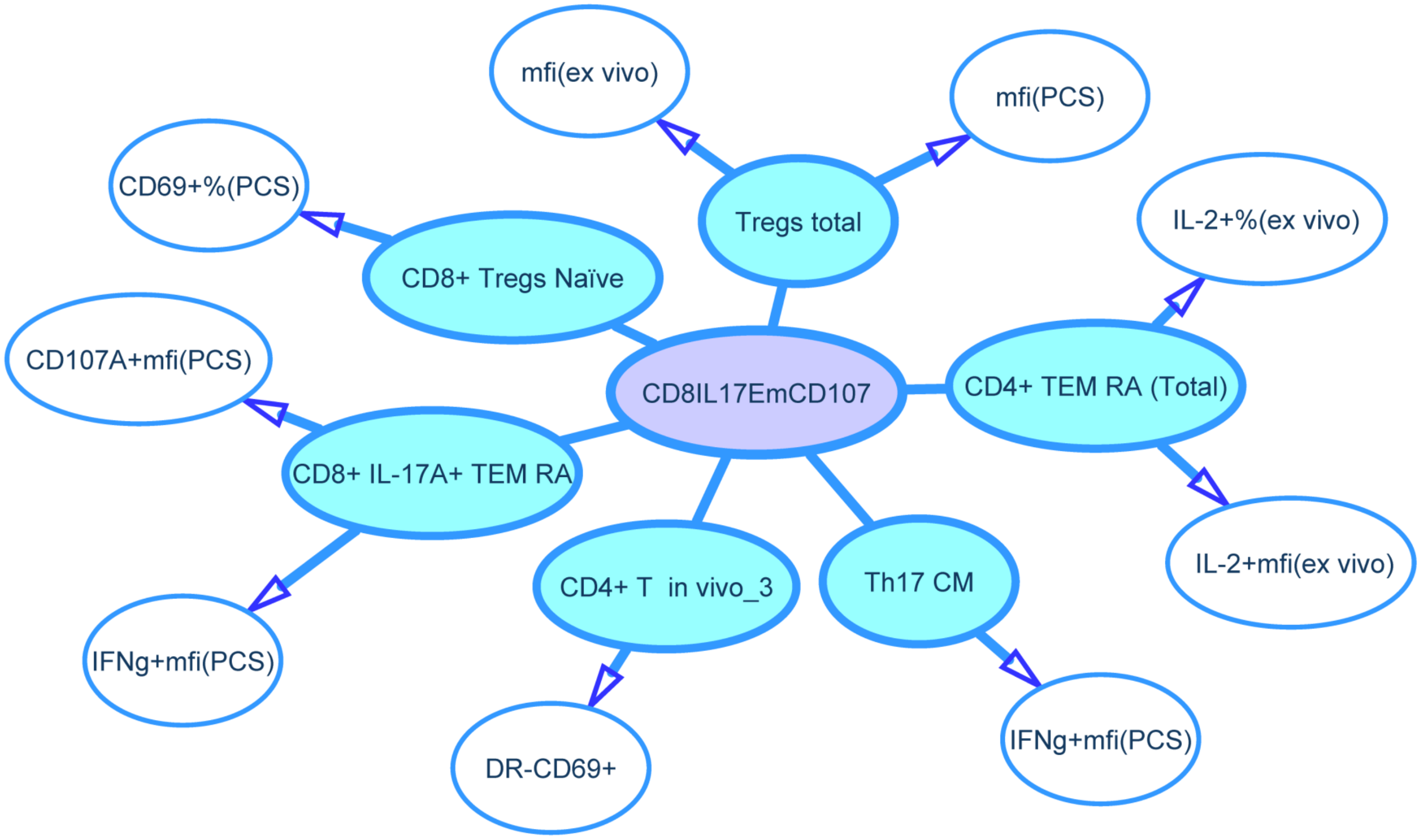


**Figure S8. Correlations of immune factors in relation to CD8IL17EmCD107.** Different T cell subsets are in blue colored circles. Uncolored circles represent either frequency or expression intensity of immune activation or cytokine markers either under ex vivo conditions or after stimulated with PCS peptides. Green colored circles indicate variants that positively correlated with vaccine efficacy by Spearman rank analysis.

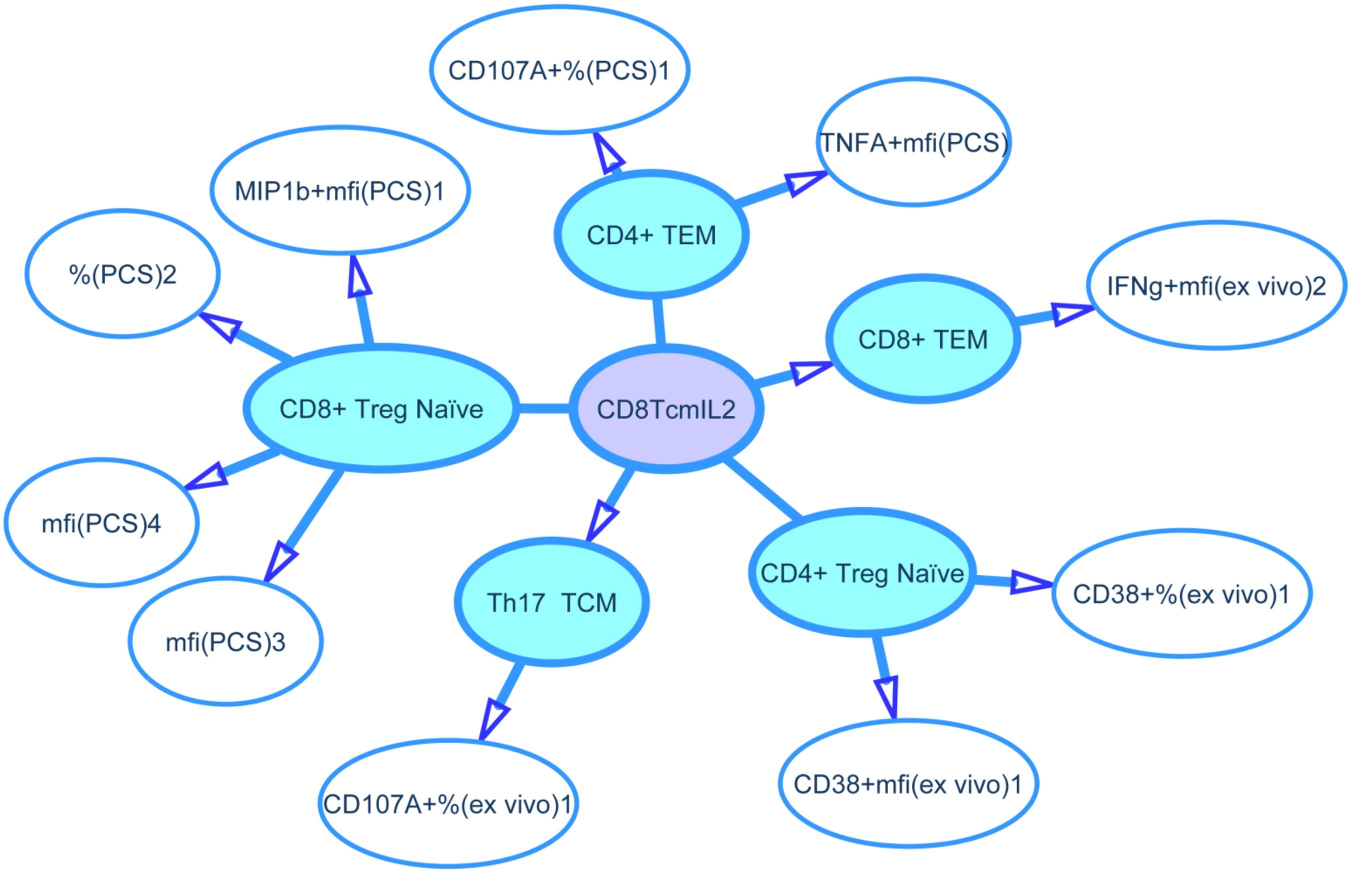


**Figure S9. Correlations of immune factors in relation to CD8TcmIL2**. Different T cell subsets are in blue colored circles. Uncolored circles represent either frequency or expression intensity of immune activation or cytokine markers either under ex vivo conditions or after stimulated with PCS peptides. Green colored circles indicate variants that positively correlated with vaccine efficacy by Spearman rank analysis.

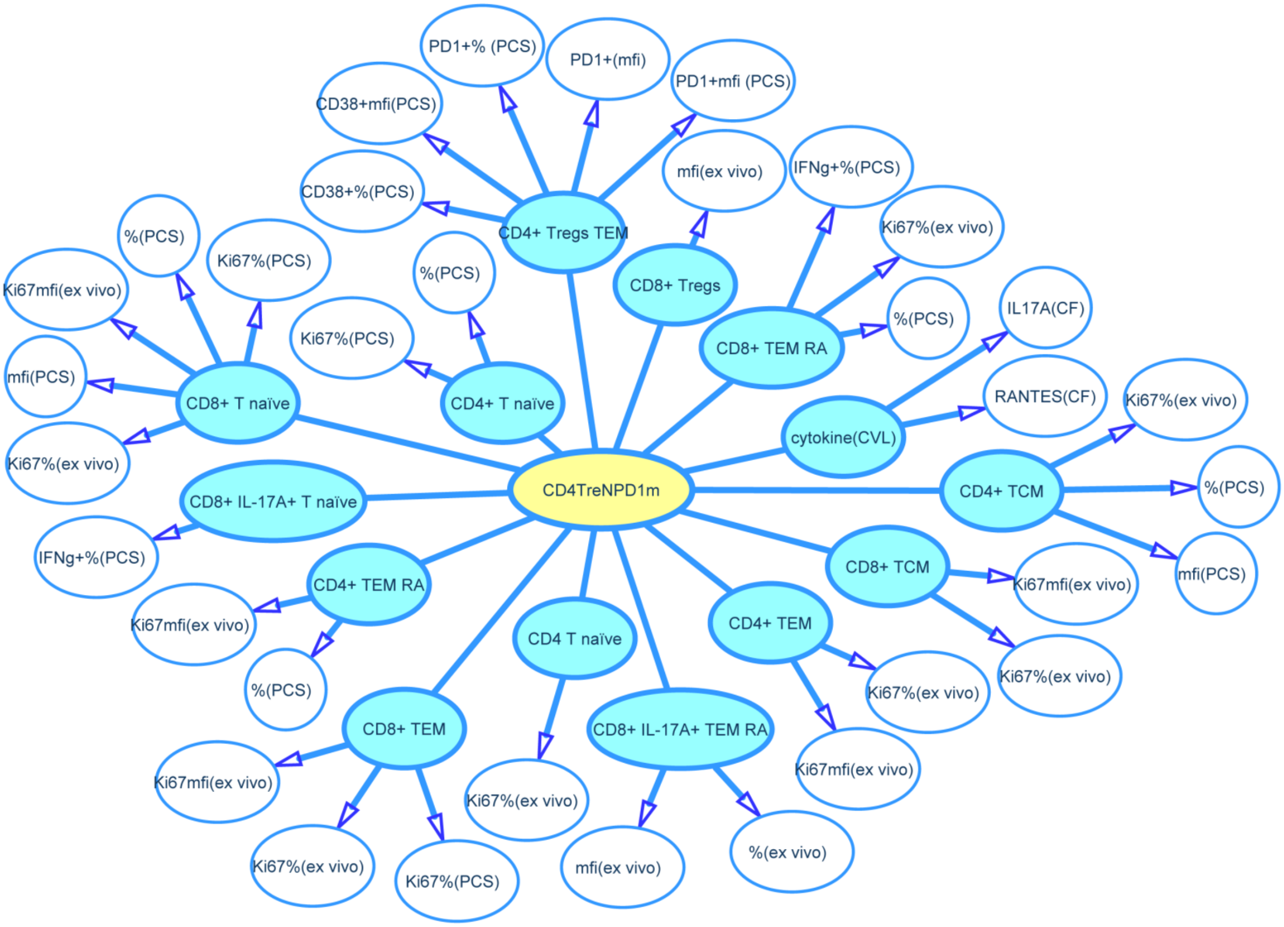


**Figure S10. Correlations of immune factors in relation to CD4TreNPD1m.** Different T cell subsets are in blue colored circles. Uncolored circles represent either frequency or expression intensity of immune activation or cytokine markers either under ex vivo conditions or after stimulated with PCS peptides. Green colored circles indicate variants that positively correlated with vaccine efficacy by Spearman rank analysis.

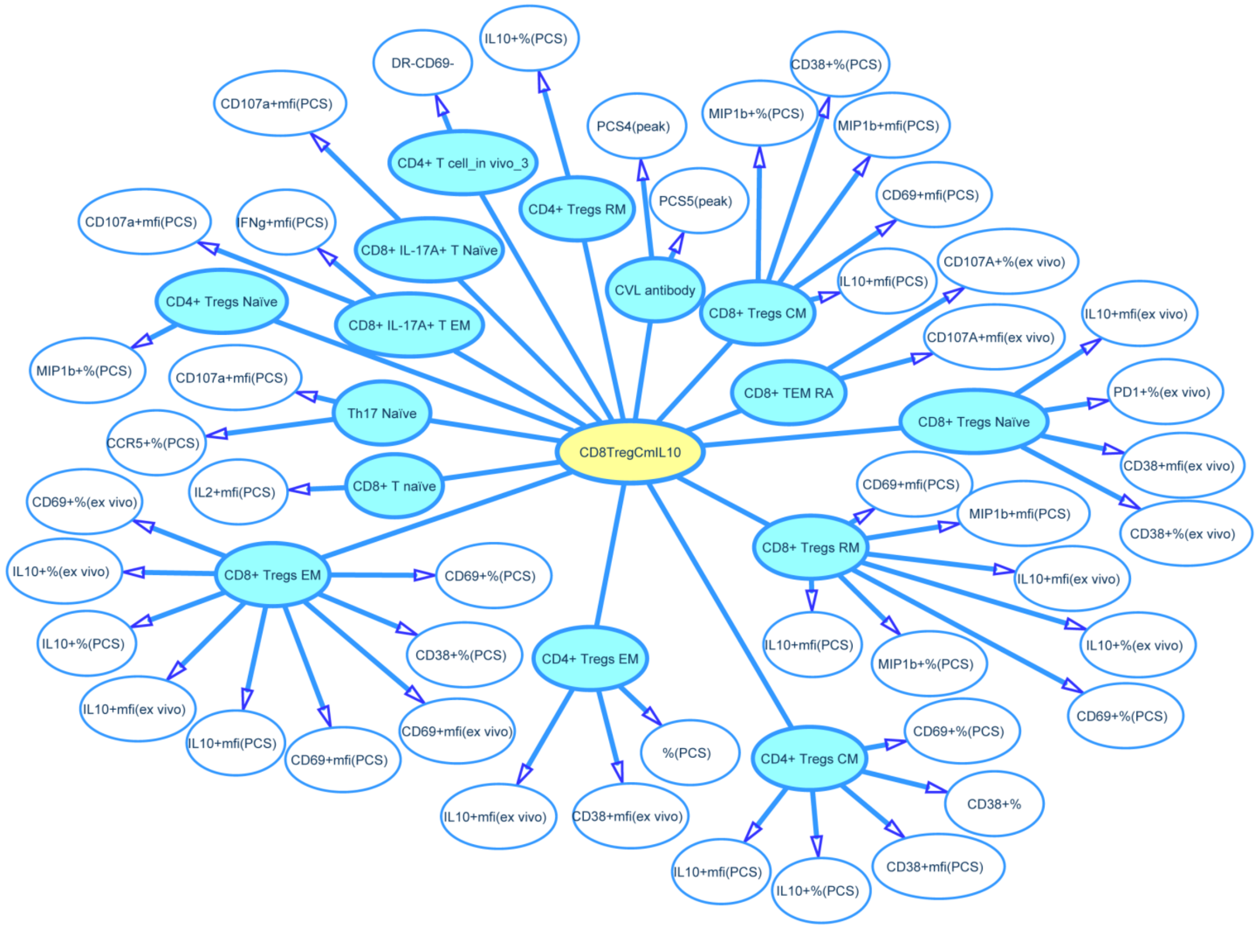


**Figure S11 Correlations of immune factors in relation to CD8TregCmIL10.** Different T cell subsets are in blue colored circles. Uncolored circles represent either frequency or expression intensity of immune activation or cytokine markers either under ex vivo conditions or after stimulated with PCS peptides. Green colored circles indicate variants that positively correlated with vaccine efficacy by Spearman rank analysis.

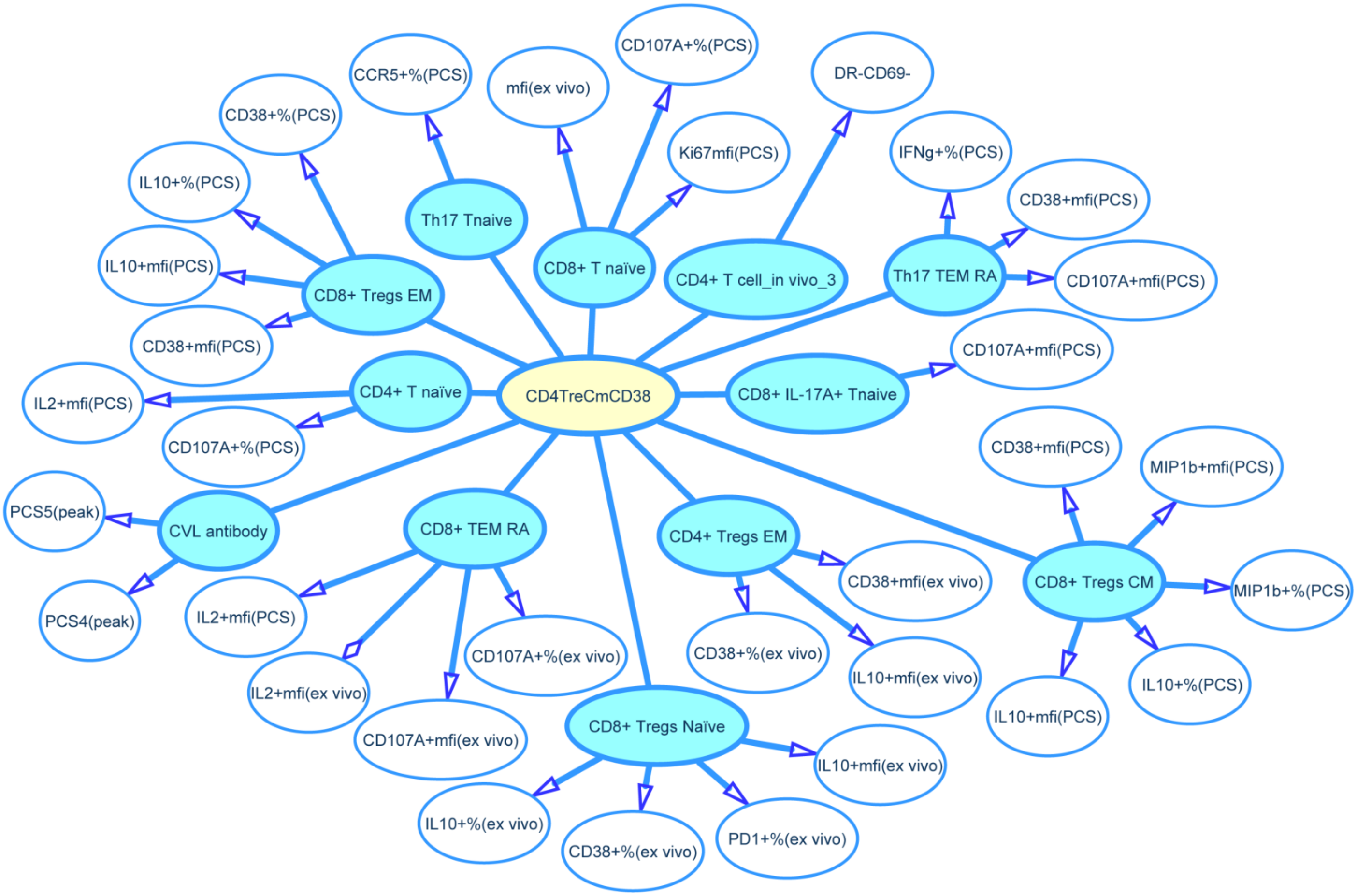


**Figure S12. Correlations of immune factors in relation to CD4TreCmCD38.** Different T cell subsets are in blue colored circles. Uncolored circles represent either frequency or expression intensity of immune activation or cytokine markers either under ex vivo conditions or after stimulated with PCS peptides. Green colored circles indicate variants that positively correlated with vaccine efficacy by Spearman rank analysis

**Table S1 Animal characteristics**

| Group | Animal ID | Gender | Age (yrs) | Weight (kgs) | MHC class I | MHC class II |
|---|---|---|---|---|---|---|
| Control | cy0774 | Female | 6 | 4.9 | M1/M4 | M1/M4 |
| Control | cy0775 | Female | 6 | 4.45 | M1/M3 | M1/M3 |
| Control | cy0776 | Female | 6 | 3.5 | M1/M2/1 | M1/M1 |
| Control | cy0777 | Female | 6 | 3.25 | M1/M1 | M1/M1 |
| Control | cy0778 | Female | 6 | 4.35 | M1/M2/1 | M1/M3 |
| Control | cy0779 | Female | 6 | 3.5 | M3/M6 | M2/M6/4/7 |
| Control | cy0780 | Female | 6 | 3.9 | M2/M2/1 | M2/M1 |
| Control | cy0781 | Female | 6 | 4.75 | M4/6/M4 | M4/6/M4 |
| PCS | cy0758 | Female | 6 | 6.65 | M6/M3/2 | M6/M2 |
| PCS | cy0759 | Female | 7 | 4.1 | M1/M4 | M1/M4 |
| PCS | cy0760 | Female | 5 | 5.85 | M1/M1 | M1/M1 |
| PCS | cy0761 | Female | 6 | 5.15 | M4/M7 | M4/M7 |
| PCS | cy0762 | Female | 7 | 4.25 | M3/M4/2 | M3/M2 |
| PCS | cy0763 | Female | 7 | 4.3 | M1/M3 | M1/M3 |
| PCS | cy0764 | Female | 6 | 7 | M1/M4 | M1/M4 |
| PCS | cy0765 | Female | 6 | 3.9 | M3/M1 | M3/M2 |

**Table S2 MHC class I and II haplotype distributions between the control and the PCS vaccine groups**

| MHC I haplotype | No. of monkeys | | P value[a] | MHC II haplotype | No. of monkeys | | P value[a] |
|---|---|---|---|---|---|---|---|
| | Control | PCS | | | Control | PCS | |
| M1 | 5 | 5 | | M1 | 6 | 4 | |
| Non-M1 | 3 | 3 | 1.0000 | Non-M1 | 2 | 4 | 0.6084 |
| M2 | 0 | 0 | | M2 | 2 | 3 | |
| Non-M2 | 8 | 8 | 1.0000 | Non-M2 | 6 | 5 | 1.0000 |
| M3 | 2 | 3 | | M3 | 2 | 3 | |
| Non-M3 | 6 | 5 | 1.0000 | Non-M3 | 6 | 5 | 1.0000 |
| M4 | 2 | 3 | | M4 | 2 | 3 | |
| Non-M4 | 6 | 5 | 1.0000 | Non-M4 | 6 | 5 | 1.0000 |
| M5 | 0 | 0 | | M5 | 0 | 0 | |
| Non-M5 | 8 | 8 | 1.0000 | Non-M5 | 8 | 8 | 1.0000 |
| M6 | 1 | 1 | | M6 | 0 | 1 | |
| Non-M6 | 7 | 7 | 1.0000 | Non-M6 | 8 | 7 | 1.0000 |
| M7 | 0 | 1 | | M7 | 0 | 1 | |
| Non-M7 | 8 | 7 | 1.0000 | Non-M7 | 8 | 7 | 1.0000 |

a. Fisher's exact test.

**Table S3 Kinetics of SIV acquistion**

| Group | Animal | Number of challenges (underlined) or days needed to acquire infection | | | | | | | | | | | | | | | | | | | | | | | | | | | | | | | | | | | | | |
|---|---|---|---|---|---|---|---|---|---|---|---|---|---|---|---|---|---|---|---|---|---|---|---|---|---|---|---|---|---|---|---|---|---|---|---|---|---|---|---|---|---|
| | | 1 | | | 2 | | | 3 | | | 4 | | | 5 | | | 6 | | | 7 | | | 8 | | | 9 | | | 10 | | | 11 | | | 12 | | | 13 | | | |
| | | 0 | 6 | 10 | 14 | 20 | 24 | 28 | 34 | 38 | 42 | 48 | 52 | 56 | 62 | 66 | 70 | 76 | 80 | 84 | 90 | 94 | 98 | 104 | 108 | 112 | 118 | 122 | 126 | 132 | 136 | 140 | 146 | 150 | 154 | 160 | 164 | 168 | 174 | 178 | 182 |
| Control | cy0774 | ○ | ○ | ○ | ○ | ○ | ○ | ● | | | | | | | | | | | | | | | | | | | | | | | | | | | | | | | | | |
| Control | cy0775 | ○ | ○ | ○ | ○ | ● | | | | | | | | | | | | | | | | | | | | | | | | | | | | | | | | | | | |
| Control | cy0776 | ○ | ● | | | | | | | | | | | | | | | | | | | | | | | | | | | | | | | | | | | | | | |
| Control | cy0777 | ○ | ○ | ○ | ○ | ○ | ○ | ○ | ○ | ○ 39 | ○ | ○ | ● 53 | | | | | | | | | | | | | | | | | | | | | | | | | | | | |
| Control | cy0778 | ○ | ○ | ○ | ○ | ○ | ○ | ○ | ○ | ○ | ○ | ○ | ○ | ○ | ○ | ○ | ○ | ○ | ○ | ○ | ○ | ○ | ○ | ○ | ○ | ○ | ○ | ○ | ○ | ○ | ○ | ○ | ○ | ○ | ○ | ○ | ○ | ○ | ○ | ○ | ○ |
| Control | cy0779 | ○ | ○ | ○ | ○ | ○ | ○ | ○ | ○ | ○ | ○ | ○ | ○ | ○ | ○ | ○ | ○ | ○ | ○ | ○ | ○ | ○ | ○ | ○ | ○ | ○ | ○ | ○ | ○ | ○ | ○ | ○ | ○ | ○ | ○ | ○ | ○ | ○ | ○ | ○ | ○ |
| Control | cy0780 | ○ | ○ | ○ | ○ | ○ | ○ | ○ | ○ | ○ | ○ | ○ | ● | | | | | | | | | | | | | | | | | | | | | | | | | | | | |
| Control | cy0781 | ○ | ○ | ○ | ○ | ● | | | | | | | | | | | | | | | | | | | | | | | | | | | | | | | | | | | |
| PCS | cy0758 | ○ | ○ | ○ | ○ | ○ | ○ | ○ | ○ | ● 39 | | | | | | | | | | | | | | | | | | | | | | | | | | | | | | | |
| PCS | cy0759 | ○ | ○ | ○ | ○ | ○ | ○ | ○ | ○ | ○ 39 | ○ | ○ | ○ 53 | ○ | ○ | ○ | ○ | ○ | ○ | ○ | ○ | ○ | ○ | ○ | ○ | ○ | ○ | ○ | ○ | ○ | ○ | ○ | ○ | ○ | ○ | ○ | ○ | ○ | ● | | |
| PCS | cy0760 | ○ | ○ | ○ | ○ | ○ | ○ | ○ | ○ | ○ 39 | ○ | ○ | ○ 53 | ○ | ○ | ○ | ○ | ○ | ○ | ○ | ○ | ○ | ○ | ○ | ○ | ○ | ○ | ○ | ○ | ○ | ○ | ○ | ○ | ○ | ○ | ○ | ○ | ○ | ○ | ○ | ○ |
| PCS | cy0761 | ○ | ○ | ○ | ○ | ○ | ○ | ○ | ○ | ○ 39 | ○ | ○ | ○ 53 | ○ | ○ | ○ | ○ | ○ | ○ | ○ | ○ | ○ | ○ | ○ | ○ | ○ | ○ | ○ | ○ | ○ | ○ | ○ | ○ | ○ | ○ | ○ | ○ | ○ | ○ | ○ | ○ |
| PCS | cy0762 | ○ | ○ | ○ | ○ | ○ | ○ | ○ | ○ | ○ | ○ | ○ | ○ | ○ | ○ | ○ | ○ | ○ | ○ | ○ | ○ | ○ | ○ | ○ | ○ | ○ | ○ | ○ | ● | | | | | | | | | | | | |
| PCS | cy0763 | ○ | ○ | ○ | ○ | ○ | ○ | ○ | ○ | ○ | ○ | ○ | ○ | ○ | ○ | ○ | ○ | ○ | ○ | ○ | ○ | ○ | ○ | ○ | ○ | ○ | ○ | ○ | ○ | ○ | ○ | ○ | ○ | ○ | ○ | ○ | ○ | ○ | ○ | ○ | ○ |
| PCS | cy0764 | ○ | ○ | ○ | ○ | ○ | ○ | ● | | | | | | | | | | | | | | | | | | | | | | | | | | | | | | | | | |
| PCS | cy0765 | ○ | ○ | ○ | ○ | ○ | ○ | ○ | ○ | ○ | ○ | ○ | ○ | ○ | ○ | ○ | ○ | ○ | ○ | ○ | ○ | ○ | ○ | ○ | ○ | ○ | ○ | ○ | ○ | ○ | ○ | ○ | ○ | ● | | | | | | | |

Intravaginal SIVmac251 challenges were conducted every two weeks. Infection status was monitored on days 0, 6, 10 and 14 of each 14-day challenge round by determining plasma viral load. Numbers of challenges (underlined) or days relative to the time of the first challenge are indicated. For logistic reason, some animals were monitored on day 39 or 53 as labelled, one day later than originally designed. Empty circle: no detectable viral load. Solid circle: positive viral load.

**Table S4. MHC class I and II haplotype distributions beteen infected and uninfected animals (Standard challenges)**

| MHC I haplotype | No. of monkeys | | P value[a] | MHC II haplotype | No. of monkeys | | P value[a] |
|---|---|---|---|---|---|---|---|
| | Infected | Uninfected | | | Infected | Uninfected | |
| M1 | 5 | 5 | | M1 | 6 | 4 | |
| Non-M1 | 3 | 3 | 1.0000 | Non-M1 | 2 | 4 | 0.6084 |
| M2 | 1 | 0 | | M2 | 2 | 3 | |
| Non-M2 | 7 | 8 | 1.0000 | Non-M2 | 6 | 5 | 1.0000 |
| M3 | 1 | 4 | | M3 | 1 | 4 | |
| Non-M3 | 7 | 4 | 0.2821 | Non-M3 | 7 | 4 | 0.2821 |
| M4 | 3 | 2 | | M4 | 3 | 2 | |
| Non-M4 | 5 | 6 | 1.0000 | Non-M4 | 5 | 6 | 1.0000 |
| M5 | 0 | 0 | | M5 | 0 | 0 | |
| Non-M5 | 8 | 8 | 1.0000 | Non-M5 | 8 | 8 | 1.0000 |
| M6 | 1 | 1 | | M6 | 1 | 0 | |
| Non-M6 | 7 | 7 | 1.0000 | Non-M6 | 7 | 8 | 1.0000 |
| M7 | 0 | 1 | | M7 | 1 | 1 | |
| Non-M7 | 8 | 7 | 1.0000 | Non-M7 | 7 | 7 | 1.0000 |

a. Fisher's exact test.

**Table S5. MHC class I and II haplotype distributions between infected and uninfected animals (extended challenges)**

| MHC I haplotype | No. of monkeys | | P value[a] | MHC II haplotype | No. of monkeys | | P value[a] |
|---|---|---|---|---|---|---|---|
| | Infected | Uninfected | | | Infected | Uninfected | |
| M1 | 7 | 3 | | M1 | 7 | 3 | |
| Non-M1 | 4 | 2 | 1.0000 | Non-M1 | 4 | 2 | 1.0000 |
| M2 | 1 | 0 | | M2 | 4 | 1 | |
| Non-M2 | 10 | 5 | 1.0000 | Non-M2 | 7 | 4 | 1.0000 |
| M3 | 3 | 2 | | M3 | 3 | 2 | |
| Non-M3 | 8 | 3 | 1.0000 | Non-M3 | 8 | 3 | 1.0000 |
| M4 | 4 | 1 | | M4 | 4 | 1 | |
| Non-M4 | 7 | 4 | 1.0000 | Non-M4 | 7 | 4 | 1.0000 |
| M5 | 0 | 0 | | M5 | 0 | 0 | |
| Non-M5 | 11 | 5 | 1.0000 | Non-M5 | 11 | 5 | 1.0000 |
| M6 | 1 | 1 | | M6 | 1 | 0 | |
| Non-M6 | 10 | 4 | 1.0000 | Non-M6 | 10 | 5 | 1.0000 |
| M7 | 0 | 1 | | M7 | 0 | 1 | |
| Non-M7 | 11 | 4 | 0.3125 | Non-M7 | 11 | 4 | 0.3125 |

a. Fisher’s exact test.

### Table S6. Cytokine-producing antigen recall responses in PBMCs from the PCS vaccine group

| Animal ID | Cytokines (pg/ml) secreted in response to PCS1-4 | | | | | | | | | | | | | | No. of challenges to cause infection |
|---|---|---|---|---|---|---|---|---|---|---|---|---|---|---|---|
| | TNF-α | IFN-γ | IL-6 | RANTES | GM-CSF | IL-1β | MCP-1 | IL-8 | MIP-1α | MIP-1β | IL-10 | IP-10 | IL-17A | IL-1α | |
| cy0758 | 0.00 | 0.00 | 1.14 | 0.00 | 0.00 | 0.00 | 0.00 | 0.99 | 0.00 | 0.00 | 0.00 | 19.51 | 0.00 | 0.00 | 3 |
| cy0759 | 43.24 | 0.00 | 0.00 | 29.47 | 0.00 | 0.00 | 0.00 | 0.00 | 0.00 | 0.00 | 0.00 | 0.00 | 0.00 | 0.00 | 13 |
| cy0760 | 0.00 | 4.84 | 3.53 | 219.59 | 0.00 | 0.00 | 0.00 | 33.47 | 0.00 | 106.80 | 1.34 | 237.04 | 0.00 | 0.00 | Uninfected |
| cy0761 | 0.00 | 0.00 | 4.44 | 943.24 | 0.00 | 0.00 | 0.00 | 0.35 | 0.00 | 0.00 | 0.00 | 0.00 | 0.00 | 0.00 | Uninfected |
| cy0762 | 0.00 | 0.00 | 0.00 | 47.85 | 0.00 | 0.00 | 0.00 | 0.00 | 0.00 | 0.00 | 0.00 | 0.00 | 0.00 | 0.00 | 9 |
| cy0763 | 0.00 | 1.13 | 22.40 | 2,498.02 | 0.00 | 0.00 | 0.00 | 8.53 | 0.00 | 563.57 | 0.00 | 0.00 | 0.00 | 0.00 | Uninfected |
| cy0764 | 0.00 | 0.00 | 0.00 | 0.00 | 0.00 | 0.00 | 0.00 | 2.37 | 0.00 | 377.49 | 0.00 | 0.00 | 0.00 | 0.00 | 2 |
| cy0765 | 0.00 | 0.00 | 0.00 | 0.00 | 19.28 | 0.00 | 0.00 | 285.19 | 0.00 | 0.00 | 0.00 | 179.60 | 0.00 | 0.00 | 11 |

| Animal ID | Cytokines (pg/ml) secreted in response to PCS5-8 | | | | | | | | | | | | | | No. of challenges to cause infection |
|---|---|---|---|---|---|---|---|---|---|---|---|---|---|---|---|
| | TNF-α | IFN-γ | IL-6 | RANTES | GM-CSF | IL-1β | MCP-1 | IL-8 | MIP-1α | MIP-1β | IL-10 | IP-10 | IL-17A | IL-1α | |
| cy0758 | 1.11 | 1.28 | 0.00 | 0.00 | 0.00 | 0.00 | 0.00 | 90.75 | 0.00 | 0.00 | 0.96 | 0.00 | 0.00 | 0.00 | 3 |
| cy0759 | 55.13 | 0.00 | 0.00 | 0.00 | 0.00 | 0.00 | 0.00 | 0.00 | 0.00 | 0.00 | 0.00 | 0.00 | 0.00 | 0.00 | 13 |
| cy0760 | 122.64 | 6.62 | 6.30 | 573.69 | 14.18 | 0.00 | 0.00 | 366.19 | 0.00 | 201.33 | 0.91 | 323.34 | 0.00 | 0.00 | Uninfected |
| cy0761 | 0.00 | 0.00 | 23.61 | 171.75 | 0.00 | 0.59 | 28.52 | 408.20 | 0.00 | 0.00 | 0.00 | 0.00 | 0.00 | 156.99 | Uninfected |
| cy0762 | 12.36 | 14.15 | 16.79 | 326.20 | 9.24 | 3.85 | 0.00 | 49.05 | 20.63 | 0.00 | 0.00 | 0.00 | 168.14 | 15.61 | 9 |
| cy0763 | 3.68 | 0.00 | 0.00 | 0.00 | 0.00 | 0.00 | 0.00 | 38.83 | 25.04 | 344.06 | 0.00 | 0.00 | 0.00 | 0.00 | Uninfected |
| cy0764 | 25.73 | 0.00 | 0.00 | 0.00 | 0.00 | 0.00 | 0.00 | 165.60 | 0.00 | 465.20 | 0.00 | 0.00 | 0.00 | 0.00 | 2 |
| cy0765 | 0.00 | 0.00 | 0.00 | 0.00 | 0.00 | 0.00 | 0.00 | 0.00 | 0.00 | 662.24 | 0.00 | 152.72 | 0.00 | 818.28 | 11 |

| Animal ID | Cytokines (pg/ml) secreted in response to PCS9-12 | | | | | | | | | | | | | | No. of challenges to cause infection |
|---|---|---|---|---|---|---|---|---|---|---|---|---|---|---|---|
| | TNF-α | IFN-γ | IL-6 | RANTES | GM-CSF | IL-1β | MCP-1 | IL-8 | MIP-1α | MIP-1β | IL-10 | IP-10 | IL-17A | IL-1α | |
| cy0758 | 0.00 | 0.80 | 0.00 | 0.00 | 0.00 | 0.00 | 0.00 | 0.00 | 0.00 | 0.00 | 0.00 | 0.00 | 0.00 | 0.00 | 3 |
| cy0759 | 0.00 | 0.00 | 0.00 | 0.00 | 0.00 | 0.00 | 0.00 | 0.00 | 0.00 | 0.00 | 0.00 | 0.00 | 0.00 | 0.00 | 13 |
| cy0760 | 0.00 | 1.02 | 47.45 | 365.24 | 0.00 | 6.90 | 62.93 | 238.46 | 56.57 | 261.44 | 0.00 | 249.40 | 0.00 | 101.05 | Uninfected |
| cy0761 | 0.00 | 0.00 | 69.48 | 205.04 | 0.00 | 7.75 | 0.00 | 76.23 | 83.67 | 312.05 | 0.00 | 0.00 | 0.00 | 0.00 | Uninfected |
| cy0762 | 0.00 | 22.46 | 14.56 | 340.55 | 15.59 | 1.51 | 0.00 | 203.23 | 1.20 | 0.00 | 8.14 | 0.00 | 142.50 | 0.00 | 9 |
| cy0763 | 28.54 | 0.00 | 12.18 | 66.59 | 0.00 | 0.00 | 0.00 | 89.41 | 33.11 | 765.01 | 0.00 | 71.05 | 0.00 | 0.00 | Uninfected |
| cy0764 | 19.58 | 0.00 | 0.00 | 0.00 | 0.00 | 0.00 | 0.00 | 0.00 | 0.00 | 126.15 | 0.00 | 0.00 | 0.00 | 0.00 | 2 |
| cy0765 | 128.23 | 0.00 | 0.00 | 0.00 | 0.00 | 0.00 | 0.00 | 0.00 | 0.00 | 773.93 | 0.00 | 143.01 | 0.00 | 69.52 | 11 |

PBMCs collected at the peak time point (week 73) were stimulated with a pool of SIV peptides (PCS1-4, PCS5-8 or PCS9-12) and quantified for cytokine secretions by Bio-Plex multiplexed cytokine assay. Number of challenges to cause infection in each animal is also listed. The values of mediium only negative control were subtracted.

**Table S7 Immune correlates of risk (CoRs)**

| Immune response | | | | Correlation with No. of challenges (Spearman rank) | |
|---|---|---|---|---|---|
| Type | Sub-type | Ag | Time point | Rho (r) | p |
| CVL mucosal cytokine | IL-1 α fold change | N/A | Peak/baseline | -0.6584 | 0.0056 |
| CVL mucosal cytokine | IL-1 α fold change | N/A | Pre-challenge/baseline | -0.5435 | 0.0296 |
| CVL mucosal cytokine | IL-1 α | N/A | Pre-challenge | -0.5168 | 0.0404 |
| CVL mucosal IgG Ab | N/A | PCS1 peptide | Peak | -0.7075 | 0.0496 |
| CVL mucosal IgG Ab | N/A | PCS3 peptide | Peak | -0.7075 | 0.0496 |
| CVL mucosal IgG Ab | N/A | PCS4 peptide | Peak | -0.7075 | 0.0496 |
| CVL mucosal IgG Ab | N/A | PCS5 peptide | Peak | -0.7807 | 0.0222 |
| CVL mucosal IgG Ab | N/A | PCS9 peptide | Peak | -0.7075 | 0.0496 |
| CD8+ T cells (ex vivo) | CCR7+CD45RA+% of CD8+ | N/A | Pre-challenge | -0.8000 | 0.0171 |
| CD4+CD25+FoxP3+ T cells (ex vivo) | CD38+% of CD4+CD25+FoxP3+CD45RA-CD127- | N/A | Pre-challenge | -0.8106 | 0.0147 |
| CD4+CD25+FoxP3+ T cells (ex vivo) | CD38 MFI of CD4+CD25+FoxP3+CD45RA-CD127- | N/A | Pre-challenge | -0.8944 | 0.0027 |
| CD8+CD25+FoxP3+ T cells (ex vivo) | CD127 MFI of CD8+CD25+FoxP3+CD45RA+ | N/A | Pre-challenge | -0.7563 | 0.0299 |
| CD8+CD25+FoxP3+ T cells (ex vivo) | CD38+% of CD8+CD25+FoxP3+CD45RA+CD127+ | N/A | Pre-challenge | -0.8665 | 0.0054 |
| CD8+CD25+FoxP3+ T cells (ex vivo) | CD38 MFI of CD8+CD25+FoxP3+CD45RA+CD127+ | N/A | Pre-challenge | -0.8665 | 0.0054 |
| CD8+CD25+FoxP3+ T cells (ex vivo) | IL-10+% of CD8+CD25+FoxP3+CD45RA+CD127+ | N/A | Pre-challenge | -0.8106 | 0.0147 |
| CD8+CD25+FoxP3+ T cells (ex vivo) | IL-10 MFI of CD8+CD25+FoxP3+CD45RA+CD127+ | N/A | Pre-challenge | -0.8665 | 0.0054 |
| CD8+CD25+FoxP3+ T cells (ex vivo) | MIP-1β+% of CD8+CD25+FoxP3+CD45RA+CD127+ | N/A | Pre-challenge | -0.8113 | 0.0145 |
| CD8+CD25+FoxP3+ T cells (ex vivo) | MIP-1β MFI of CD8+CD25+FoxP3+CD45RA+CD127+ | N/A | Pre-challenge | -0.8318 | 0.0104 |
| CD8+CD25+FoxP3+ T cells (ex vivo) | PD-1+% of CD8+CD25+FoxP3+CD45RA+CD127+ | N/A | Pre-challenge | -0.8665 | 0.0054 |
| CD8+CD25+FoxP3+ T cells (ex vivo) | PD-1 MFI of CD8+CD25+FoxP3+CD45RA+CD127+ | N/A | Pre-challenge | -0.8106 | 0.0147 |
| CD4+IL17A+ T cells (Ag recall) | CCR5+% of CD4+IL17A+CCR7+CD45RA+ | PCS1-12 peptide pool | Pre-challenge | -0.7507 | 0.0319 |
| CD8+IL17A+ T cells (Ag recall) | CCR5+% of CD8+IL17A+CCR7+CD45RA+ | PCS1-12 peptide pool | Pre-challenge | -0.7278 | 0.0407 |
| CD8+IL17A+ T cells (Ag recall) | CD69+% of CD8+IL17A+CCR7+CD45RA+ | PCS1-12 peptide pool | Pre-challenge | -0.7278 | 0.0407 |
| CD8+IL17A+ T cells (Ag recall) | CD69 MFI of CD8+IL17A+CCR7+CD45RA+ | PCS1-12 peptide pool | Pre-challenge | -0.7278 | 0.0407 |
| CD8+IL17A+ T cells (Ag recall) | CD107a MFI of CD8+IL17A+CCR7+CD45RA+ | PCS1-12 peptide pool | Pre-challenge | -0.7507 | 0.0319 |
| CD8+IL17A+ T cells (Ag recall) | IFN- γ MFI of CD8+IL17A+CCR7-CD45RA- | PCS1-12 peptide pool | Pre-challenge | -0.7507 | 0.0319 |
| CD4+CD25+FoxP3+ T cells (Ag recall) | MIP-1 β+% of CD4+CD25+FoxP3+CD45RA+CD127+ | PCS1-12 peptide pool | Pre-challenge | -0.7507 | 0.0319 |
| CD4+CD25+FoxP3+ T cells (Ag recall) | CD38+% of CD4+CD25+FoxP3+CD45RA-CD127+ | PCS1-12 peptide pool | Pre-challenge | -0.7507 | 0.0319 |
| CD4+CD25+FoxP3+ T cells (Ag recall) | CD38 MFI of CD4+CD25+FoxP3+CD45RA-CD127+ | PCS1-12 peptide pool | Pre-challenge | -0.7826 | 0.0217 |
| CD8+CD25+FoxP3+ T cells (Ag recall) | CD38+% of CD8+CD25+FoxP3+CD45RA-CD127+ | PCS1-12 peptide pool | Pre-challenge | -0.8665 | 0.0054 |
| CD8+CD25+FoxP3+ T cells (Ag recall) | CD38 MFI of CD8+CD25+FoxP3+CD45RA-CD127+ | PCS1-12 peptide pool | Pre-challenge | -0.8106 | 0.0147 |
| CD8+CD25+FoxP3+ T cells (Ag recall) | IL-10 MFI of CD8+CD25+FoxP3+CD45RA-CD127+ | PCS1-12 peptide pool | Pre-challenge | -0.7826 | 0.0217 |
| CD8+CD25+FoxP3+ T cells (Ag recall) | MIP-1β MFI of CD8+CD25+FoxP3+CD45RA-CD127+ | PCS1-12 peptide pool | Pre-challenge | -0.7826 | 0.0217 |
| CD8+CD25+FoxP3+ T cells (Ag recall) | CD38+% of CD8+CD25+FoxP3+CD45RA-CD127- | PCS1-12 peptide pool | Pre-challenge | -0.7507 | 0.0319 |
| CD8+CD25+FoxP3+ T cells (Ag recall) | IL-10+% of CD8+CD25+FoxP3+CD45RA-CD127- | PCS1-12 peptide pool | Pre-challenge | -0.7507 | 0.0319 |
| CD8+CD25+FoxP3+ T cells (Ag recall) | IL-10 MFI of CD8+CD25+FoxP3+CD45RA-CD127- | PCS1-12 peptide pool | Pre-challenge | -0.7507 | 0.0319 |

Immune factors significantly correlating with number of challenges needed to cause infection were determined by Spearman rank correlation analysis. Ag: antigen (used to quantify antibody or to stimulate cells). CVL: cervicovaginal lavage wash. N/A: not applicable. Peak time point: Week 73 of vaccination schedule (1 week after final vaccination boost). Pre-challenge time point: Week 90 of vaccination schedule. Ab: antibody. Ex vivo: peripheral blood mononuclear cells (PBMCs) were not stimulated with Ag. Ag recall: PBMCs were stimulated with Ag in culture. MFI: mean fluorescent intensity. All analyses were conducted in the PCS vaccine group (n=8), except that CVL mucosal cytoine analyses were conducted in the PCS vaccine and control groups combined (n=16).

**Table S8. The index of immune variants positively correlated with 6 features in the LASSO model**

| Featrue name | Feature variant | variant category | variant | Spearman Rho | P-value |
|---|---|---|---|---|---|
| CD4TregEmPD1 | CD4+ Tregs EM PD1+% | Tcell_cytokine | IL6 (PCS1-4) | 0.900 | 0.002 |
| CD4TregEmPD1 | CD4+ Tregs EM PD1+% | Tcell_cytokine | IL6 (PCS9-12) | 0.780 | 0.022 |
| CD4TregEmPD1 | CD4+ Tregs EM PD1+% | Tcell_cytokine | RANTES(PCS1-4) | 0.866 | 0.005 |
| CD4TregEmPD1 | CD4+ Tregs EM PD1+% | Tcell_cytokine | MIP1a(PCS9-12) | 0.900 | 0.002 |
| CD4TregEmPD1 | CD4+ Tregs EM PD1+% | Tcell_cytokine | MIP1b(PCS_Pr) | 0.738 | 0.037 |
| CD4TregEmPD1 | CD4+ Tregs EM PD1+% | Th17 EM | %(ex vivo) | 0.866 | 0.005 |
| CD4TregEmPD1 | CD4+ Tregs EM PD1+% | CD8+ IL-17A+ CM | %(ex vivo) | 0.845 | 0.008 |
| CD4TregEmPD1 | CD4+ Tregs EM PD1+% | CD8+ IL-17A+ EM | %(ex vivo) | 0.845 | 0.008 |
| CD4TregEmPD1 | CD4+ Tregs EM PD1+% | CD4+ Tregs RM | %(PCS) | 0.866 | 0.005 |
| CD4TregEmPD1 | CD4+ Tregs EM PD1+% | CD4+ Tregs RM | mfi(PCS) | 0.751 | 0.032 |
| CD4TregEmPD1 | CD4+ Tregs EM PD1+% | CD4+ Tregs RM | CD38+%(PCS) | 0.926 | 0.001 |
| CD4TregEmPD1 | CD4+ Tregs EM PD1+% | CD4+ Tregs RM | CD38+mfi(PCS) | 0.780 | 0.022 |
| CD4TregEmPD1 | CD4+ Tregs EM PD1+% | CD8+ T naïve | IFNg+mfi(ex vivo) | 0.794 | 0.019 |
| CD4TregEmPD1 | CD4+ Tregs EM PD1+% | CD8+ T in vivo_3 | DR+CD69- | 0.866 | 0.012 |
| CD8IL17EmCD107 | CD8+ IL17A+ TEM CD107a+% | CD4+ T in vivo_3 | DR-CD69+ | 0.793 | 0.034 |
| CD8IL17EmCD107 | CD8+ IL17A+ TEM CD107a+% | CD8+ IL-17A+ TEM RA | CD107A+mfi(PCS) | 0.956 | 0.000 |
| CD8IL17EmCD107 | CD8+ IL17A+ TEM CD107a+% | CD8+ IL-17A+ TEM RA | IFNg+mfi(PCS) | 0.852 | 0.007 |
| CD8IL17EmCD107 | CD8+ IL17A+ TEM CD107a+% | Th17 CM | IFNg+mfi(PCS) | 0.906 | 0.002 |
| CD8IL17EmCD107 | CD8+ IL17A+ TEM CD107a+% | Tregs total | mfi(ex vivo) | 0.945 | 0.000 |
| CD8IL17EmCD107 | CD8+ IL17A+ TEM CD107a+% | Tregs total | mfi(PCS) | 0.737 | 0.037 |
| CD8IL17EmCD107 | CD8+ IL17A+ TEM CD107a+% | CD8+ Tregs Naïve | CD69+%(PCS) | 0.802 | 0.017 |
| CD8IL17EmCD107 | CD8+ IL17A+ TEM CD107a+% | CD4+ TEM RA (Total) | IL-2+%(ex vivo) | 0.766 | 0.027 |
| CD8IL17EmCD107 | CD8+ IL17A+ TEM CD107a+% | CD4+ TEM RA (Total) | IL-2+mfi(ex vivo) | 0.796 | 0.032 |
| CD8TcmIL2 | CD8+TCM IL2+% | Th17 TCM | CD107A+%(ex vivo) | 0.730 | 0.040 |
| CD8TcmIL2 | CD8+TCM IL2+% | CD4+ Treg Naïve | CD38+%(ex vivo) | 0.788 | 0.020 |
| CD8TcmIL2 | CD8+TCM IL2+% | CD4+ Treg Naïve | CD38+mfi(ex vivo) | 0.788 | 0.020 |
| CD8TcmIL2 | CD8+TCM IL2+% | CD8+ Treg Naïve | %(PCS) | 0.752 | 0.031 |
| CD8TcmIL2 | CD8+TCM IL2+% | CD8+ Treg Naïve | mfi(PCS) | 0.792 | 0.019 |
| CD8TcmIL2 | CD8+TCM IL2+% | CD8+ Treg Naïve | MIP1b+mfi(PCS) | 0.739 | 0.036 |
| CD8TcmIL2 | CD8+TCM IL2+% | CD8+ Treg Naïve | mfi(PCS) | 0.788 | 0.020 |
| CD8TcmIL2 | CD8+TCM IL2+% | CD8+ TEM | IFNg+mfi(ex vivo) | 0.803 | 0.016 |
| CD8TcmIL2 | CD8+TCM IL2+% | CD4+ TEM | CD107A+%(PCS) | 0.873 | 0.005 |
| CD8TcmIL2 | CD8+TCM IL2+% | CD4+ TEM | TNFA+mfi(PCS) | 0.755 | 0.030 |
| CD4TreNPD1m | CD4+Treg Naïve PD1+ mfi | cytokine(CVL) | RANTES(CF) | 0.710 | 0.048 |
| CD4TreNPD1m | CD4+Treg Naïve PD1+ mfi | cytokine(CVL) | IL17A(CF) | 0.710 | 0.048 |
| CD4TreNPD1m | CD4+Treg Naïve PD1+ mfi | CD8+ IL-17A+ TEM RA | %(ex vivo) | 0.759 | 0.029 |
| CD4TreNPD1m | CD4+Treg Naïve PD1+ mfi | CD8+ IL-17A+ TEM RA | mfi(ex vivo) | 0.741 | 0.036 |
| CD4TreNPD1m | CD4+Treg Naïve PD1+ mfi | CD8+ IL-17A+ T naïve | IFNg+%(PCS) | 0.757 | 0.030 |
| CD4TreNPD1m | CD4+Treg Naïve PD1+ mfi | CD8+ Tregs | mfi(ex vivo) | 0.888 | 0.003 |
| CD4TreNPD1m | CD4+Treg Naïve PD1+ mfi | CD4+ Tregs TEM | PD1+(mfi) | 0.843 | 0.009 |
| CD4TreNPD1m | CD4+Treg Naïve PD1+ mfi | CD4+ Tregs TEM | CD38+%(PCS) | 0.814 | 0.014 |
| CD4TreNPD1m | CD4+Treg Naïve PD1+ mfi | CD4+ Tregs TEM | CD38+mfi(PCS) | 0.814 | 0.014 |
| CD4TreNPD1m | CD4+Treg Naïve PD1+ mfi | CD4+ Tregs TEM | PD1+% (PCS) | 0.843 | 0.009 |
| CD4TreNPD1m | CD4+Treg Naïve PD1+ mfi | CD4+ Tregs TEM | PD1+mfi (PCS) | 0.814 | 0.014 |
| CD4TreNPD1m | CD4+Treg Naïve PD1+ mfi | CD4 T naïve | Ki67%(ex vivo) | 0.823 | 0.023 |
| CD4TreNPD1m | CD4+Treg Naïve PD1+ mfi | CD4+ TEM RA | Ki67mfi(ex vivo) | 0.823 | 0.023 |
| CD4TreNPD1m | CD4+Treg Naïve PD1+ mfi | CD4+ TCM | Ki67%(ex vivo) | 0.823 | 0.023 |
| CD4TreNPD1m | CD4+Treg Naïve PD1+ mfi | CD4+ TEM | Ki67%(ex vivo) | 0.823 | 0.023 |
| CD4TreNPD1m | CD4+Treg Naïve PD1+ mfi | CD4+ TEM | Ki67mfi(ex vivo) | 0.898 | 0.006 |
| CD4TreNPD1m | CD4+Treg Naïve PD1+ mfi | CD8+ T naïve | Ki67%(ex vivo) | 0.823 | 0.023 |
| CD4TreNPD1m | CD4+Treg Naïve PD1+ mfi | CD8+ T naïve | Ki67mfi(ex vivo) | 0.823 | 0.023 |
| CD4TreNPD1m | CD4+Treg Naïve PD1+ mfi | CD8+ TEM RA | Ki67%(ex vivo) | 0.898 | 0.006 |
| CD4TreNPD1m | CD4+Treg Naïve PD1+ mfi | CD8+ TCM | Ki67%(ex vivo) | 0.898 | 0.006 |
| CD4TreNPD1m | CD4+Treg Naïve PD1+ mfi | CD8+ TCM | Ki67mfi(ex vivo) | 0.898 | 0.006 |
| CD4TreNPD1m | CD4+Treg Naïve PD1+ mfi | CD8+ TEM | Ki67%(ex vivo) | 0.898 | 0.006 |
| CD4TreNPD1m | CD4+Treg Naïve PD1+ mfi | CD8+ TEM | Ki67mfi(ex vivo) | 0.898 | 0.006 |
| CD4TreNPD1m | CD4+Treg Naïve PD1+ mfi | CD4+ T naïve | %(PCS) | 0.759 | 0.029 |
| CD4TreNPD1m | CD4+Treg Naïve PD1+ mfi | CD4+ T naïve | Ki67%(PCS) | 0.759 | 0.029 |

**Table S8. The index of immune variants positively correlated with the 6 features in the LASSO model (continue)**

| | | | | | |
|---|---|---|---|---|---|
| CD4TreNPD1m | CD4+Treg Naïve PD1+ mfi | CD4+ TEM RA | %(PCS) | 0.715 | 0.046 |
| CD4TreNPD1m | CD4+Treg Naïve PD1+ mfi | CD4+ TCM | %(PCS) | 0.811 | 0.015 |
| CD4TreNPD1m | CD4+Treg Naïve PD1+ mfi | CD4+ TCM | mfi(PCS) | 0.952 | 0.000 |
| CD4TreNPD1m | CD4+Treg Naïve PD1+ mfi | CD8+ T naïve | %(PCS) | 0.719 | 0.044 |
| CD4TreNPD1m | CD4+Treg Naïve PD1+ mfi | CD8+ T naïve | mfi(PCS) | 0.952 | 0.000 |
| CD4TreNPD1m | CD4+Treg Naïve PD1+ mfi | CD8+ T naïve | Ki67%(PCS) | 0.820 | 0.013 |
| CD4TreNPD1m | CD4+Treg Naïve PD1+ mfi | CD8+ TEM RA | %(PCS) | 0.719 | 0.044 |
| CD4TreNPD1m | CD4+Treg Naïve PD1+ mfi | CD8+ TEM RA | IFNg+%(PCS) | 0.916 | 0.001 |
| CD4TreNPD1m | CD4+Treg Naïve PD1+ mfi | CD8+ TEM | Ki67%(PCS) | 0.804 | 0.016 |
| CD8TregCmIL10 | CD8+ Tregs CM IL10+% | CVL antibody | PCS4(peak) | 0.764 | 0.027 |
| CD8TregCmIL10 | CD8+ Tregs CM IL10+% | CVL antibody | PCS5(peak) | 0.733 | 0.039 |
| CD8TregCmIL10 | CD8+ Tregs CM IL10+% | CD4+ Tregs CM | CD38+% | 0.959 | 0.000 |
| CD8TregCmIL10 | CD8+ Tregs CM IL10+% | CD4+ T cell_in vivo_3 | DR-CD69- | 0.794 | 0.033 |
| CD8TregCmIL10 | CD8+ Tregs CM IL10+% | Th17 Naïve | CCR5+%(PCS) | 0.959 | 0.000 |
| CD8TregCmIL10 | CD8+ Tregs CM IL10+% | Th17 Naïve | CD107a+mfi(PCS) | 0.756 | 0.030 |
| CD8TregCmIL10 | CD8+ Tregs CM IL10+% | CD8+ IL-17A+ T Naïve | CD107a+mfi(PCS) | 0.959 | 0.000 |
| CD8TregCmIL10 | CD8+ Tregs CM IL10+% | CD8+ IL-17A+ T EM | CD107a+mfi(PCS) | 0.756 | 0.030 |
| CD8TregCmIL10 | CD8+ Tregs CM IL10+% | CD8+ IL-17A+ T EM | IFNg+mfi(PCS) | 0.959 | 0.000 |
| CD8TregCmIL10 | CD8+ Tregs CM IL10+% | CD4+ Tregs EM | CD38+mfi(ex vivo) | 0.875 | 0.004 |
| CD8TregCmIL10 | CD8+ Tregs CM IL10+% | CD4+ Tregs EM | IL10+mfi(ex vivo) | 0.751 | 0.032 |
| CD8TregCmIL10 | CD8+ Tregs CM IL10+% | CD8+ Tregs Naïve | CD38+%(ex vivo) | 0.839 | 0.009 |
| CD8TregCmIL10 | CD8+ Tregs CM IL10+% | CD8+ Tregs Naïve | CD38+mfi(ex vivo) | 0.750 | 0.032 |
| CD8TregCmIL10 | CD8+ Tregs CM IL10+% | CD8+ Tregs Naïve | IL10+mfi(ex vivo) | 0.839 | 0.009 |
| CD8TregCmIL10 | CD8+ Tregs CM IL10+% | CD8+ Tregs Naïve | PD1+%(ex vivo) | 0.839 | 0.009 |
| CD8TregCmIL10 | CD8+ Tregs CM IL10+% | CD8+ Tregs RM | IL10+%(ex vivo) | 0.756 | 0.030 |
| CD8TregCmIL10 | CD8+ Tregs CM IL10+% | CD8+ Tregs RM | IL10+mfi(ex vivo) | 0.756 | 0.030 |
| CD8TregCmIL10 | CD8+ Tregs CM IL10+% | CD8+ Tregs EM | CD69+%(ex vivo) | 0.756 | 0.030 |
| CD8TregCmIL10 | CD8+ Tregs CM IL10+% | CD8+ Tregs EM | CD69+mfi(ex vivo) | 0.756 | 0.030 |
| CD8TregCmIL10 | CD8+ Tregs CM IL10+% | CD8+ Tregs EM | IL10+%(ex vivo) | 0.756 | 0.030 |
| CD8TregCmIL10 | CD8+ Tregs CM IL10+% | CD8+ Tregs EM | IL10+mfi(ex vivo) | 0.756 | 0.030 |
| CD8TregCmIL10 | CD8+ Tregs CM IL10+% | CD4+ Tregs Naïve | MIP1b+%(PCS) | 0.959 | 0.000 |
| CD8TregCmIL10 | CD8+ Tregs CM IL10+% | CD4+ Tregs RM | IL10+%(PCS) | 0.756 | 0.030 |
| CD8TregCmIL10 | CD8+ Tregs CM IL10+% | CD4+ Tregs CM | CD38+mfi(PCS) | 1.000 | . |
| CD8TregCmIL10 | CD8+ Tregs CM IL10+% | CD4+ Tregs CM | CD69+%(PCS) | 0.756 | 0.030 |
| CD8TregCmIL10 | CD8+ Tregs CM IL10+% | CD4+ Tregs CM | IL10+%(PCS) | 0.839 | 0.009 |
| CD8TregCmIL10 | CD8+ Tregs CM IL10+% | CD4+ Tregs CM | IL10+mfi(PCS) | 0.875 | 0.004 |
| CD8TregCmIL10 | CD8+ Tregs CM IL10+% | CD4+ Tregs EM | %(PCS) | 0.756 | 0.030 |
| CD8TregCmIL10 | CD8+ Tregs CM IL10+% | CD8+ Tregs RM | CD69+%(PCS) | 0.756 | 0.030 |
| CD8TregCmIL10 | CD8+ Tregs CM IL10+% | CD8+ Tregs RM | CD69+mfi(PCS) | 0.756 | 0.030 |
| CD8TregCmIL10 | CD8+ Tregs CM IL10+% | CD8+ Tregs RM | IL10+mfi(PCS) | 0.756 | 0.030 |
| CD8TregCmIL10 | CD8+ Tregs CM IL10+% | CD8+ Tregs RM | MIP1b+%(PCS) | 0.756 | 0.030 |
| CD8TregCmIL10 | CD8+ Tregs CM IL10+% | CD8+ Tregs RM | MIP1b+mfi(PCS) | 0.756 | 0.030 |
| CD8TregCmIL10 | CD8+ Tregs CM IL10+% | CD8+ Tregs CM | CD38+%(PCS) | 0.839 | 0.009 |
| CD8TregCmIL10 | CD8+ Tregs CM IL10+% | CD8+ Tregs CM | CD69+mfi(PCS) | 0.756 | 0.030 |
| CD8TregCmIL10 | CD8+ Tregs CM IL10+% | CD8+ Tregs CM | IL10+mfi(PCS) | 1.000 | . |
| CD8TregCmIL10 | CD8+ Tregs CM IL10+% | CD8+ Tregs CM | MIP1b+%(PCS) | 0.875 | 0.004 |
| CD8TregCmIL10 | CD8+ Tregs CM IL10+% | CD8+ Tregs CM | MIP1b+mfi(PCS) | 1.000 | . |
| CD8TregCmIL10 | CD8+ Tregs CM IL10+% | CD8+ Tregs EM | CD38+%(PCS) | 0.959 | 0.000 |
| CD8TregCmIL10 | CD8+ Tregs CM IL10+% | CD8+ Tregs EM | CD69+%(PCS) | 0.756 | 0.030 |
| CD8TregCmIL10 | CD8+ Tregs CM IL10+% | CD8+ Tregs EM | CD69+mfi(PCS) | 0.756 | 0.030 |
| CD8TregCmIL10 | CD8+ Tregs CM IL10+% | CD8+ Tregs EM | IL10+%(PCS) | 0.959 | 0.000 |
| CD8TregCmIL10 | CD8+ Tregs CM IL10+% | CD8+ Tregs EM | IL10+mfi(PCS) | 0.959 | 0.000 |
| CD8TregCmIL10 | CD8+ Tregs CM IL10+% | CD8+ TEM RA | CD107A+%(ex vivo) | 0.765 | 0.027 |
| CD8TregCmIL10 | CD8+ Tregs CM IL10+% | CD8+ TEM RA | CD107A+mfi(ex vivo) | 0.765 | 0.027 |
| CD8TregCmIL10 | CD8+ Tregs CM IL10+% | CD8+ T naïve | IL2+mfi(PCS) | 0.711 | 0.048 |
| CD4TreCmCD38 | CD4+ Tregs CM CD38+% | CD8+ Tregs CM | IL10+%(PCS) | 0.959 | 0.000 |
| CD4TreCmCD38 | CD4+ Tregs CM CD38+% | CVL antibody | PCS4(peak) | 0.733 | 0.039 |
| CD4TreCmCD38 | CD4+ Tregs CM CD38+% | CVL antibody | PCS5(peak) | 0.764 | 0.027 |
| CD4TreCmCD38 | CD4+ Tregs CM CD38+% | CD4+ T cell_in vivo_3 | DR-CD69- | 0.794 | 0.033 |
| CD4TreCmCD38 | CD4+ Tregs CM CD38+% | Th17 Tnaive | CCR5+%(PCS) | 1.000 | . |
| CD4TreCmCD38 | CD4+ Tregs CM CD38+% | Th17 TEM RA | CD38+mfi(PCS) | 0.750 | 0.032 |
| CD4TreCmCD38 | CD4+ Tregs CM CD38+% | Th17 TEM RA | CD107A+mfi(PCS) | 0.750 | 0.032 |

**Table S8. The index of immune variants positively correlated with 6 features in the LASSO model (continue)**

| | | | | | |
|---|---|---|---|---|---|
| CD4TreCmCD38 | CD4+ Tregs CM CD38+% | Th17 TEM RA | IFNg+%(PCS) | 0.750 | 0.032 |
| CD4TreCmCD38 | CD4+ Tregs CM CD38+% | CD8+ IL-17A+ Tnaive | CD107A+mfi(PCS) | 1.000 | . |
| CD4TreCmCD38 | CD4+ Tregs CM CD38+% | CD4+ Tregs EM | CD38+%(ex vivo) | 0.750 | 0.032 |
| CD4TreCmCD38 | CD4+ Tregs CM CD38+% | CD4+ Tregs EM | CD38+mfi(ex vivo) | 0.839 | 0.009 |
| CD4TreCmCD38 | CD4+ Tregs CM CD38+% | CD4+ Tregs EM | IL10+mfi(ex vivo) | 0.783 | 0.022 |
| CD4TreCmCD38 | CD4+ Tregs CM CD38+% | CD8+ Tregs Naïve | CD38+%(ex vivo) | 0.875 | 0.004 |
| CD4TreCmCD38 | CD4+ Tregs CM CD38+% | CD8+ Tregs Naïve | IL10+%(ex vivo) | 0.750 | 0.032 |
| CD4TreCmCD38 | CD4+ Tregs CM CD38+% | CD8+ Tregs Naïve | IL10+mfi(ex vivo) | 0.875 | 0.004 |
| CD4TreCmCD38 | CD4+ Tregs CM CD38+% | CD8+ Tregs Naïve | PD1+%(ex vivo) | 0.875 | 0.004 |
| CD4TreCmCD38 | CD4+ Tregs CM CD38+% | CD8+ Tregs CM | CD38+mfi(PCS) | 0.750 | 0.032 |
| CD4TreCmCD38 | CD4+ Tregs CM CD38+% | CD8+ Tregs CM | IL10+mfi(PCS) | 0.959 | 0.000 |
| CD4TreCmCD38 | CD4+ Tregs CM CD38+% | CD8+ Tregs CM | MIP1b+%(PCS) | 0.839 | 0.009 |
| CD4TreCmCD38 | CD4+ Tregs CM CD38+% | CD8+ Tregs CM | MIP1b+mfi(PCS) | 0.959 | 0.000 |
| CD4TreCmCD38 | CD4+ Tregs CM CD38+% | CD8+ Tregs EM | CD38+%(PCS) | 1.000 | . |
| CD4TreCmCD38 | CD4+ Tregs CM CD38+% | CD8+ Tregs EM | CD38+mfi(PCS) | 0.756 | 0.030 |
| CD4TreCmCD38 | CD4+ Tregs CM CD38+% | CD8+ Tregs EM | IL10+%(PCS) | 1.000 | . |
| CD4TreCmCD38 | CD4+ Tregs CM CD38+% | CD8+ Tregs EM | IL10+mfi(PCS) | 1.000 | . |
| CD4TreCmCD38 | CD4+ Tregs CM CD38+% | CD8+ T naïve | mfi(ex vivo) | 0.713 | 0.047 |
| CD4TreCmCD38 | CD4+ Tregs CM CD38+% | CD8+ TEM RA | CD107A+%(ex vivo) | 0.819 | 0.013 |
| CD4TreCmCD38 | CD4+ Tregs CM CD38+% | CD8+ TEM RA | CD107A+mfi(ex vivo) | 0.819 | 0.013 |
| CD4TreCmCD38 | CD4+ Tregs CM CD38+% | CD8+ TEM RA | IL2+mfi(ex vivo) | 0.770 | 0.043 |
| CD4TreCmCD38 | CD4+ Tregs CM CD38+% | CD4+ T naïve | CD107A+%(PCS) | 0.756 | 0.030 |
| CD4TreCmCD38 | CD4+ Tregs CM CD38+% | CD4+ T naïve | IL2+mfi(PCS) | 0.761 | 0.028 |
| CD4TreCmCD38 | CD4+ Tregs CM CD38+% | CD8+ T naïve | CD107A+%(PCS) | 0.756 | 0.030 |
| CD4TreCmCD38 | CD4+ Tregs CM CD38+% | CD8+ T naïve | Ki67mfi(PCS) | 0.756 | 0.030 |
| CD4TreCmCD38 | CD4+ Tregs CM CD38+% | CD8+ TEM RA | IL2+mfi(PCS) | 0.756 | 0.030 |

### Table S9. A List of capture and detection antibodies, and protein standards used in the Bio-Plex Multiplexed cytokine/chemokine assays

| **Protein Standards** | | |
|---|---|---|
| **Name** | **Catalog No.** | **Manufacturer** |
| Recombinant Human CXCL8/IL-8 Protein | 208-IL-10 | R&D system |
| Recombinant Human CCL5/RANTES Protein | 278-RN-10 | R&D system |
| Recombinant Human GM-CSF Protein | 215-GM | R&D system |
| Recombinant Human Interferon Gamma Protein | RIFNG50 | ThemoFisher scientific |
| Recombinant Human IL-1 beta/IL-1F2 | 201-LB | R&D system |
| Recombinant Human IL-6 Protein | 206-IL-10 | R&D system |
| Recombinant Human IL-10 (aa 19-178) Protein | 1064-IL | R&D system |
| Recombinant Human IL-17A Protein | 317-ILB | R&D system |
| Recombinant Human CXCL10/IP-10 | 266-IP | R&D system |
| Recombinant Human CCL2/MCP-1 Protein | 279-MC | R&D system |
| Recombinant Human CCL3/MIP-1 alpha protein | 270-LD-10 | R&D system |
| Recombinant Human CCL4/MIP-1 beta Protein | 271-BME-10 | R&D system |
| Recombinant Human TNF-alpha Protein | 210-TA-20 | R&D system |
| Recombinant Human IL-1 alpha | 200-LA | R&D system |
| **Capture Antibodies** | | |
| **Name** | **Catalog No.** | **Manufacturer** |
| Human CXCL8/IL-8 MAb | M801 | ThermoFisher Scientific |
| Human CCL5/RANTES PAb | P230E | ThermoFisher Scientific |
| Rat Anti-Human GM-CSF-UNLB | 10111-01 | SouthernBiotech |
| Human IFNγ MAb | M700A | ThermoFisher Scientific |
| Human IL-1beta/ IL-1F2 Antibody | MAB601-500 | R&D System |
| Human IL-6 MAb | M620 | ThermoFisher Scientific |
| Rat Anti-Human IL-10-UNLB | 10100-01 | SouthernBiotech |
| Human/Primate IL-17/IL-17A Antibody | MAB317-500 | R&D System |
| Human IP-10/ CXCL10/CRG-2 Antibody | MAB266-500 | R&D System |
| Human MCP-1/CCL2/JE Antibody | MAB679-500 | R&D System |
| Human MIP-1α/CCL3Antibody | AF-270-NA | R&D System |
| Human MIP-1β/CCL4 Antibody | MAB271-100 | R&D System |
| Human TNFα MAb | M303 | ThermoFisher Scientific |
| Human IL-1 alpha/IL-1F1 Mab | MAB200 | R&D system |
| **Detection Antibodies** | | |
| **Name** | **Catalog No.** | **Manufacturer** |
| Human CXCL8/IL-8 MAb, Biotin-labeled | M802B | ThermoFisher Scientific |
| Human CCL5/RANTES MAb, Biotin-labeled | M230B | ThermoFisher Scientific |
| Rat Anti-Human GM-CSF-BIOT | 10112-08 | SouthernBiotech |
| Human IFNγ MAb, Biotin-labeled | M701B | ThermoFisher Scientific |
| Human IL-1beta/IL-1F2 Biotinylated Antibody | BAF201 | R&D System |
| Human IL-6 MAb, Biotin-labeled | M621B | ThermoFisher Scientific |
| Rat Anti-Human IL-10-BIOT | 10110-08 | SouthernBiotech |
| Human/Primate IL-17/IL-17A Biotinylated Antibody | BAF317 | R&D System |
| Human IP-10/CXCL10/CRG-2 Biotinylated Antibody | BAF266 | R&D System |
| Human MCP-1/CCL2/JE Biotinylated Antibody | BAF279 | R&D System |
| Human MIP-1α/CCL3 Biotinylated Antibody | BAF270 | R&D System |
| Human MIP-1β/CCL4 Biotinylated Antibody | BAF271 | R&D System |
| Human TNFα MAb, Biotin-labeled | M302B | ThermoFisher Scientific |
| Human IL-1 alpha/IL-1F1 Biotinylated | BAF200 | R&D system |

**Table S10 Final concentration of the standard pool in the Bio-Plex Multiplexed cytokine/chemokine assays**

| Name | Concentration in pg/mL |
|---|---|
| Recombinant Human CXCL8/IL-8 Protein | 27000 |
| Recombinant Human CCL5/RANTES Protein | 18500 |
| Recombinant Human GM-CSF Protein | 17000 |
| Recombinant Human Interferon Gamma Protein | 10500 |
| Recombinant Human IL-1 beta/IL-1F2 | 19500 |
| Recombinant Human IL-6 Protein | 9500 |
| Recombinant Human IL-10 (aa 19-178) Protein | 28000 |
| Recombinant Human IL-17A Protein | 38000 |
| Recombinant Human CXCL10/IP-10 | 24000 |
| Recombinant Human CCL2/MCP-1 Protein | 26500 |
| Recombinant Human CCL3/MIP-1 alpha protein | 27500 |
| Recombinant Human CCL4/MIP-1 beta Protein | 9000 |
| Recombinant Human TNF-alpha Protein | 18000 |
| Recombinant Human IL-1 alpha | 160000 |

**Table S11. SIV PCS peptides used to stimulate PBMCs for testing antigen recall responses**

| Peptide | Peptide Sequence |
|---|---|
| PCS1-1 | [H]APSSGRGGNYPVQQI[OH] |
| PCS1-2 | [H]SGRGGNYPVQQIGGN[OH] |
| PCS1-3 | [H]RGGNYPVQQIGGNYV[OH] |
| PCS2-1 | [H]GGPGQKARLMAEALK[OH] |
| PCS2-2 | [H]GQKARLMAEALKEAL[OH] |
| PCS2-3 | [H]KARLMAEALKEALAP[OH] |
| PCS3-1 | [H]LAPVPIPFAAAQQRG[OH] |
| PCS3-2 | [H]VPIPFAAAQQRGPRK[OH] |
| PCS3-3 | [H]IPFAAAQQRGPRKPI[OH] |
| PCS4-1 | [H]MAKCPDRQAGFLGLG[OH] |
| PCS4-2 | [H]CPDRQAGFLGLGPWG[OH] |
| PCS4-3 | [H]DRQAGFLGLGPWGKK[OH] |
| PCS5-1 | [H]GPWGKKPRNFPMAQVHQGLM[OH] |
| PCS5-2 | [H]YGQMPRQTGGFFRPWSMGKE[OH] |
| PCS5-3 | [H]KPRNFPMAQVHQGLM[OH] |
| PCS6-1 | [H]YGQMPRQTGGFFRPW[OH] |
| PCS6-2 | [H]MPRQTGGFFRPWSMG[OH] |
| PCS6-3 | [H]RQTGGFFRPWSMGKE[OH] |
| PCS7-1 | [H]WSMGKEAPQFPHGSS[OH] |
| PCS7-2 | [H]GKEAPQFPHGSSASG[OH] |
| PCS7-3 | [H]EAPQFPHGSSASGAD[OH] |
| PCS8-1 | [H]LQGGDRGFAAPQFSL[OH] |
| PCS8-2 | [H]GDRGFAAPQFSLWRR[OH] |
| PCS8-3 | [H]RGFAAPQFSLWRRPV[OH] |
| PCS9-1 | [H]LTALGMSLNFPIAKV[OH] |
| PCS9-2 | [H]LGMSLNFPIAKVEPV[OH] |
| PCS9-3 | [H]MSLNFPIAKVEPVKV[OH] |
| PCS10-1 | [H]KDPIEGEETYYTDGS[OH] |
| PCS10-2 | [H]IEGEETYYTDGSCNK[OH] |
| PCS10-3 | [H]GEETYYTDGSCNKQS[OH] |
| PCS11-1 | [H]LVSQGIRQVLFLEKI[OH] |
| PCS11-2 | [H]QGIRQVLFLEKIEPA[OH] |
| PCS11-3 | [H]IRQVLFLEKIEPAQE[OH] |
| PCS12-1 | [H]NQGQYMNTPWRNPAE[OH] |
| PCS12-2 | [H]QYMNTPWRNPAEERE[OH] |
| PCS12-3 | [H]MNTPWRNPAEEREKL[OH] |

### Table S12. Flow cytometry panel design

| T cell Panel 1 | | |
|---|---|---|
| Marker | **Fluorochrome** | **Clone** |
| Live/Dead | Blue (405) | reactive dye |
| CD3 | V500 | SP34-2 |
| CD4 | BV605 | L200 |
| CD8 | V450 | RP8-T4 |
| CCR7 | A700 | 150503 |
| CD127 (IL-7) | PE-CF594 | HIL-7R-M2 |
| CD107a | PE-Cy5 | H4:A3 |
| IL-2 | FITC | MQ1-17H12 |
| IFNγ | BV786 | 4S.B3 |
| TNFα | APC | MAb11 |
| Ki67 | PE-Cy7 | B56 |
| MIP1b | PE | D21-1351 |
| CD45RO | APC-H7 | UCHL1 |
| IL-17A | BV650 | N49-653 |
| Marker | **Fluorochrome** | **Clone** |
| **T cell Panel 2** | | |
| Marker | **Fluorochrome** | **Clone** |
| Live/Dead | Blue (405) | reactive dye |
| CD3 | V500 | SP34-2 |
| CD4 | BV605 | L200 |
| CD8 | APC-Cy7 | RP8-T4 |
| CCR7 | Alexa700 | 150503 |
| CD38 | FITC | HB7 |
| PD-1 | BV650 | M1H4 |
| CD69 | APC | FN50 |
| Foxp3 | V450 | 259D/C7 |
| CD127 | Pe-CF594 | HIL-7R-M2 |
| IL-10 | BV711 | JES3-9D7 |
| CD25 | PE-CY5 | M-A251 |
| CD45RA | PE-Cy7 | L48 |
| MIP1b | PE | D21-1351 |
| **Live/Dead** | **Blue (405)** | **reactive dye** |
| CD3 | V500 | SP34-2 |
| CD4 | BV605 | L200 |
| CD8 | V450 | RP8-T4 |
| CCR7 | A700 | 150503 |
| CD127 (IL-7) | PE-CF594 | HIL-7R-M2 |
| MIP1b | PE | D21-1351 |
| CD45RO | APC-H7 | UCHL1 |
| IL-17A | BV650 | N49-653 |
| CD195 (CCR5) | PE-Cy7 | 2D7/CCR5 |
| CD38 | FITC | HB7 |
| CD69 | APC | FN50 |
| CD107a | PE-Cy5 | H4:A3 |
| IFNγ | BV786 | 4S.B3 |
